# Pore-C sequencing identifies episome-driven chromosome conformation perturbations differentiating pneumococcal epigenetic variants

**DOI:** 10.1101/2025.03.18.643841

**Authors:** Tze Yee Lim, Samuel T. Horsfield, Catherine M. Troman, Stephen D. Bentley, Min Jung Kwun, Nicholas J. Croucher

**Affiliations:** MRC Centre for Global Infectious Disease Analysis, Department of Infectious Disease Epidemiology, School of Public Health, Imperial College London, London, W12 0BZ, UK; European Molecular Biology Laboratory, European Bioinformatics Institute, Wellcome Genome Campus, Hinxton, CB10 1SA, UK; Parasites & Microbes, Wellcome Sanger Institute, Wellcome Genome Campus, Hinxton, Cambridge, CB10 1SA, UK

## Abstract

*Streptococcus pneumoniae* (pneumococcus) is a genetically diverse opportunistic bacterial pathogen that expresses two phase-variable loci encoding restriction-modification systems. Comparisons of two genetically-distinct pairs of epigenetically-distinct variants, each distinguished by a stabilised arrangement of one of these phase-variable loci, found the consequent changes in genome-wide DNA methylation patterns were associated with differential expression of mobile genetic elements (MGEs). This relationship was hypothesised to be mediated through changes in xenogenic silencing (XS) or nucleoid organisation. Therefore the chromosomal conformation of the both variants of each isolate were characterised using Illumina Hi-C, and Nanopore Pore-C, sequencing. Both methods concurred that the organisation of the pneumococcal chromosome was dominated by small-scale structures, with most pairwise interactions between loci <25 kb apart. Neither found substantial evidence for higher-order structure or XS in the pneumococcal genome, with more complex contact patterns only evident around the replication origin. Comparisons between the variants identified phage-related chromosomal islands (PRCIs) as the foci of differential contact densities between the variants. This was driven by copy number variation, resulting from variable excision and replication of the episomal PRCIs. However, the methods were discordant in their identification of the variant in which the PRCI was more actively replicating in both pairs. Validatory experiments demonstrated that the prevalence of circular PRCIs was not determined by DNA modification, but instead varied stochastically between colonies in both backgrounds, and was metastable during vegetative growth. PRCI excision was inducible by mitomycin C, but independent of the presence of a phage. Yet transcriptional activation of these elements was affected by both signals, indicating transcription and replication are separately regulated. Therefore pneumococcal MGEs do not appear to be subject to XS, resulting in heterogeneity being generated within these bacterial populations through the frequent local disruption of chromosome conformation resulting from the stochastic excision and reintegration of episomal elements.

**Author summary:** The pneumococcus is a bacterium with a circular chromosome that is organised by DNA-binding proteins and often contains mobile genetic elements (MGEs), genes able to transmit between bacteria. All pneumococci encode defences against MGEs, some of which create epigenetic modifications (typically methylation) genome-wide at particular sequence motifs. Changes in these epigenetic patterns are associated with altered MGE gene expression and replication in otherwise genetically-identical bacteria. To test whether this was the result of methylation remodelling the organisation of the chromosome, we compared the contact patterns across the chromosome using two sequencing technologies. Both methods concurred that the pneumococcal genome is generally folded into small structures, with the biggest differences between the variants caused by the replication of MGEs. However, the methods disagreed on the variant in which the MGEs replicated fastest. Further experiments showed that MGE replication was stable over the course of culturing over hours, but would randomly change level between days, explaining the inconsistent observations. MGE replication was found to rise in response to DNA damage, whereas gene expression also depended on the presence of other signals, explaining the discrepancies in these activities between variants. Hence MGEs significantly contribute to the heterogeneity that rapidly accumulates within pneumococcal populations.

## Introduction

*Streptococcus pneumoniae* (the pneumococcus) is a gram-positive commensal bacterium and opportunistic pathogen. Pneumococci are a common cause of diseases such as conjunctivitis, pneumonia, sepsis and meningitis [1]. The species has an extensive pangenome [2–4] as a consequence of exchange through transformation [5] and the frequent integration of mobile genetic elements (MGEs) into the pneumococcal chromosome [4]. However, the acquisition of genes can be inhibited by restriction modification systems (RMSs) [6,7]. Almost all pneumococci share two phase-variable Type I RMSs: the *Spn*III RMS encoded by the inverting variable restriction (*ivr*) locus [8,9], and the *Spn*IV RMS encoded by the translocating variable restriction (*tvr*) locus [4,9]. The rapid variation of these loci can inhibit the spread of prophage within clonally-related populations [10,11]. Yet the changes also cause global alterations to the methylation of the genome. Hence the sensitivity of pneumococcal transcription to epigenomic changes means these RMSs also cause phenotypic heterogeneity between isolates [9,12].

Variation in the *Spn*III RMS has been associated with changes in capsule expression [9,13,14], human carriage [15], adhesion to human cells [16], and virulence in a mouse model [9,17]. Variation at the *tvr* locus is instead associated with variation in transformation efficiency [12,18]. However, it has not been possible to identify methylation sites that can explain the transcriptional changes through proximal effects at gene regulatory loci [9,12]. Yet methylation has broader effects on the biophysical properties of DNA that means a global change in methylation patterns could affect the conformation of the bacterial nucleoid, and alter the interactions between regulatory proteins and DNA loci [19]. Hence it is plausible that the epigenetic effects are instead mediated through longer-distance, larger-scale changes in chromosomal conformation.

This is supported by analysis of two pneumococcal isolates, RMV7 and RMV8. Mutations disrupting the recombinase that catalyses rearrangements at the *tvr* locus enabled “locked” phase variants of each to be isolated: one representing the common, dominant arrangement (RMV7_domi_ and RMV8_domi_), and one expressing a rare arrangement (RMV7_rare_ and RMV8_rare_) [7]. These variants differ in the methylation motif targeted by the *Spn*IV RMS, but are otherwise isogenic outside the *tvr* loci. Both rare variants were more transformable than their dominant variant counterparts, and in both cases this was at least partly attributable to the altered activity of MGEs integrated into the chromosome [12,18]. RMV7 contains a cryptic integrative and conjugative element next to *rpsI* (ICE*_rpsI_*) and the antibiotic-resistance transposon Tn*916*, as well as two phage-related chromosomal islands (PRCIs; alternatively known as phage-inducible chromosomal islands, or PICIs) integrated next to *dnaN* (PRCI*_dnaN_*) and *uvrA* (PRCI*_uvrA_*). PRCI*_dnaN_* was more transcriptionally active in RMV7_domi_, suppressing the induction of competence for transformation through inducing stress responses [12]. RMV8 encodes a prophage (ϕRMV8), a PRCI integrated within the *mal* operon (PRCI*_malA_*), and a CIPhR prophage remnant (previously annotated as PRCI*_tadA_*) [18,20]. In RMV8_domi_, reduced excision of ϕRMV8 resulted in greater expression of a prophage-modified RNA that suppressed competence induction [18]. However, no specific methylation sites have been identified that could explain this difference in MGE activity [12].

Hence host DNA conformation is a plausible mechanism by which epigenetic modification could modulate the activity of MGEs through xenogeneic silencing (XS) [21]. This process depends on nucleoid-associated proteins (NAPs) that preferentially bind to the AT-rich DNA that is characteristic of MGEs [22,23], and are capable of silencing their gene expression. As well as affecting MGE gene transcription [23–25], these NAPs can modulate the excision and re-integration of MGEs [26]. Such differences in XS activity could explain the inter-variant differences observed in RMV7 and RMV8. At least four families of NAPs have been identified [21,27–30]. Two are common in gram-negative *Proteobacteria*: H-NS, identified in *Escherichia coli* [31], and MvaT, characterised in *Pseudomonas* [30]. By contrast, Lsr2 has only been found in gram-positive *Actinobacteria* [29], and the Rok protein has only been characterised in *Bacillus subtilis* [32]. Hence no XS system has yet been characterised in *S. pneumoniae*. Nevertheless, the species encodes orthologues of Rok, as well as other proteins that preferentially bind AT-rich DNA [33]. Therefore the differential activity of MGEs in RMV7 and RMV8 may be attributable to changes in the patterns of NAPs associated with the genome.

The conformation of the genome, and therefore the distribution of NAPs and regions subject to XS, can be characterised using chromosome conformation capture (3C) [34]. This approach typically involves crosslinking DNA to proteins with formaldehyde, digesting the DNA with a frequently-cutting restriction enzyme, then facilitating proximity ligation to link DNA fragments that were spatially close within the cell (Fig. 1A). The subsequent use of paired-end Illumina data enables such contact patterns to be assayed at a genome-wide scale using high-throughput 3C (Hi-C), which infers instances of digestion and re-ligation within the DNA separating the two reads based on their mapping positions and orientations relative to a reference genome [35]. More recently, long-read data generated by Nanopore sequencing has enabled 3C analyses to use information from concatemers consisting of multiple DNA fragments ligated together [36,37]. Hence this Pore-C method makes it possible to infer higher-order multiway chromatin interactions, and represents an efficient method for generating data on the scale of a bacterial genome.

**Figure 1.**
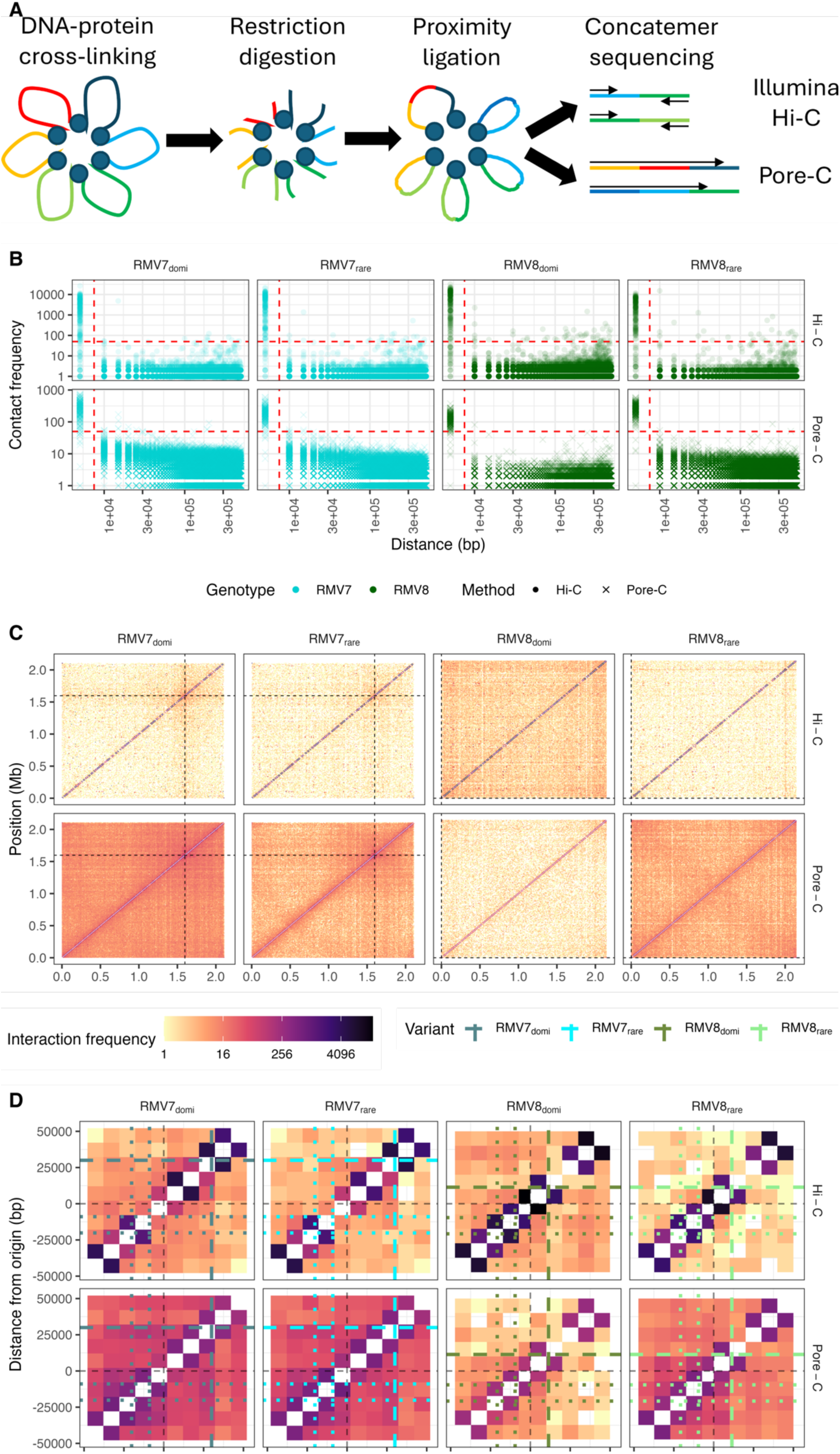
Chromosomal conformation analysis of the epigenetic variants of two pneumococcal genotypes. **(A)** Diagram describing the methodology underpinning the use of high-throughput sequencing to capture chromatin conformation (3C). The final sequencing step distinguishes the Illumina (Hi-C) and Oxford Nanopore Technology (Pore-C) methods. **(B)** Relationship between the frequency of contacts between loci and their separation in the genome. Each panel represents a unique combination of sequencing technology and epigenetic variant. Each point corresponds to a detected pairwise interaction between loci in a contact matrix calculated at a 5 kb resolution for both technologies. The red dashed lines represent the thresholds used to define the closely-spaced loci between which particularly high contact densities were inferred. **(C)** Genome-wide Hi-C and Pore-C contact matrices shown at a 10 kb resolution. Each panel displays the cumulative number of contacts identified in three biological replicates for each unique combination of sequencing technology and epigenetic variant. The frequency of detected interactions between loci across the genome is represented by the colour of the corresponding cell in these symmetrical matrices. The black dashed lines indicate the position of the origin of replication. **(D)** Matrix detailing the cumulative number of contacts identified in loci surrounding the origin of replication, the position of which is marked by black dashed lines. The coloured dashed lines indicate the position of the *parS* site downstream of *ori*; the dotted lines show the position of the *parS* sites upstream of *ori*. Each cell in the matrix represents data from a 10 kb locus.

Therefore this study compared Hi-C and Pore-C approaches to inferring the chromosomal conformation of the dominant and rare variants of *S. pneumoniae* RMV7 and RMV8. These assays were employed to determine whether it is feasible that the phenotypic heterogeneity associated with epigenetic variation can be attributed to changes in XS, or other alterations in the distribution of NAPs across the genome.

## Results

### Short-range interactions dominate pneumococcal Hi-C and Pore-C data

Three independent 3C preparations were generated for each of *S. pneumoniae* RMV7_domi_, RMV7_rare_, RMV8_domi_ and RMV8_rare_ (Table S1, S2). These were digested with *Nla*III, which can cut at sites found in the pneumococcal chromosome at intervals of ∼300 bp (Table S3), and used to generate a 12-plex library for Nanopore sequencing. However, this yielded few reads, as the activity of the pores declined rapidly during the sequencing run (Fig. S1). Residual DNA from these samples was instead processed for Illumina sequencing, and successfully generated Hi-C data (Table S2). A further three independent 3C preparations were generated for each of the four variants using *Mlu*CI, which can cut at sites found in the pneumococcal chromosome at intervals of ∼125 bp (Table S3). Previous applications of the Pore-C approach to human cells found an additional protease treatment with pronase increased the efficacy of de-cross-linking, thereby slowing the rate of nanopore blocking by proteins attached to DNA [38]. Correspondingly, including this step in the sample preparation protocol enabled more efficient data generation from individual flow cells (Fig. S1; Table S2).

The three biological replicates for both methods for each of the variants were mapped to the appropriate dominant variant reference genome (Fig. S2, S3; Table S1). The necessity of mapping short read fragments meant the Pore-C data, which contains a higher rate of sequencing errors, mapped less efficiently than the Hi-C data. Nevertheless, a high proportion of the Nanopore data could be uniquely mapped to locations within the pneumococcal genome (Fig. S3). To establish the highest resolution at which analyses could be reliably conducted, the proportions of inferred relative mapping orientations of sequence fragments were plotted against the separation between their mapped locations if they came from the same read pair, for Illumina Hi-C data, or from the same read, for Pore-C data. This found that sequence fragments separated by less than 5 kb were typically mapped to opposite strands of the genome for Illumina Hi-C data (Fig. S4), or to the same strand of the genome, for Pore-C data (Fig. S5). These are the orientations expected for undigested DNA sequenced with these technologies, consistent with only a subset of restriction sites being cleaved during sample preparation. This is typical of 3C preparations, requiring a convergence distance to be defined as the separation at which sufficient digestion and re-ligation occurred to ensure all sequence fragment mapping orientations become equally frequent [39]. This was estimated to be between 5 and 10 kb for both technologies (Fig. S4-5).

Plotting the overall frequency of pairs against their separation highlighted particularly strong interactions at an interval of ∼5 kb (Fig. 1B), which may indicate a particular set of atypical interactions. However, plotting the locations of these high-level short-range interactions found they were distributed across the chromosome, and therefore may represent false positive interactions resulting from incompletely-digested DNA (Fig. S6). Therefore data were subsequently analysed at a resolution of 10 kb, to mitigate any effects of DNA fragments that had not undergone digestion and religation.

This locus size was used to analyse the contact frequencies between all loci across the chromosome for each replicate (Fig. S7-8), and combined across each set of replicates (Fig. 1C). A strong diagonal was evident across all of the symmetrical contact frequency matrices. This suggested most interactions occurred over distances <25 kb, consistent with a simple genome organisation, dominated by short-range structures. There were relatively few signals of higher-level chromosomal organisation, apart from an elevated density of off-diagonal interactions evident around the origin of replication (*ori*; Fig. 1D, S8). This was positioned ∼1.6 Mb from the beginning of the RMV7 reference, resulting in vertical and horizontal stripes across the matrix (Fig. 1C). The RMV8 *ori* was at the start of the reference genome, resulting instead in a framing effect, with elevated intensities in each corner of the matrix.

Focussing on the origin of replication suggested the presence of boundaries to interactions that were consistent across technologies at ∼40 kb or ∼20 kb downstream of *ori* in RMV7 and RMV8, respectively (Fig. 1D). These coincide with the location of the only conserved *parS* site in this downstream replichore [40], which is more distant from *ori* in RMV7 as a consequence of the integration of PRCI*_dnaN_* in the intervening sequence [41]. The *parS* site is recognised by the ParB chromosome segregation machinery, which is integral to organising the origin domain of the chromosomes of the Bacillota species *B. subtilis* [42] and *S. pneumoniae* [40]. Hence this motif is likely to underlie the observed disruption to chromosomal contacts in this region. Although two *parS* sites are proximal to *ori* in the upstream replichore, these do not appear to affect contact patterns in the same way, consistent with previous observations of *parS* sites having distinct effects depending on their location [40,42]. Hence both Hi-C and Pore-C data produced a consistent view of the contacts that dominate the pneumococcal genome’s conformation.

### Differential interactions between variants span PRCIs

A genome-wide search for differences in chromosome conformation between the variants was undertaken using contact matrices that summarised contact counts within non-overlapping 10 bp loci. The DiffHiC (Fig. S9-S12) and multiHiCcompare (Fig. S13-S16) methods were used, both of which fit generalised linear models to independent biological replicate datasets to identify differences in contact frequencies [43,44]. However, they differ in their normalisation of the data [45]. Clustering of the biological replicates by the diffHiC pipeline suggested the variants could be resolved from one another by the genome-wide patterns of contacts in each comparison, except in the Hi-C analysis of the RMV7 variants (Fig. S17). However, these separations were not always associated with high bootstrap values, suggesting the differences were attributable to specific loci, rather than broad changes in genome-wide contact frequencies. Correspondingly, specific sites exhibiting significantly different contact frequencies between the genotypes, following Benjamini-Hochberg corrections for multiple testing, could be identified using Manhattan plots (Fig. 2A).

**Figure 2.**
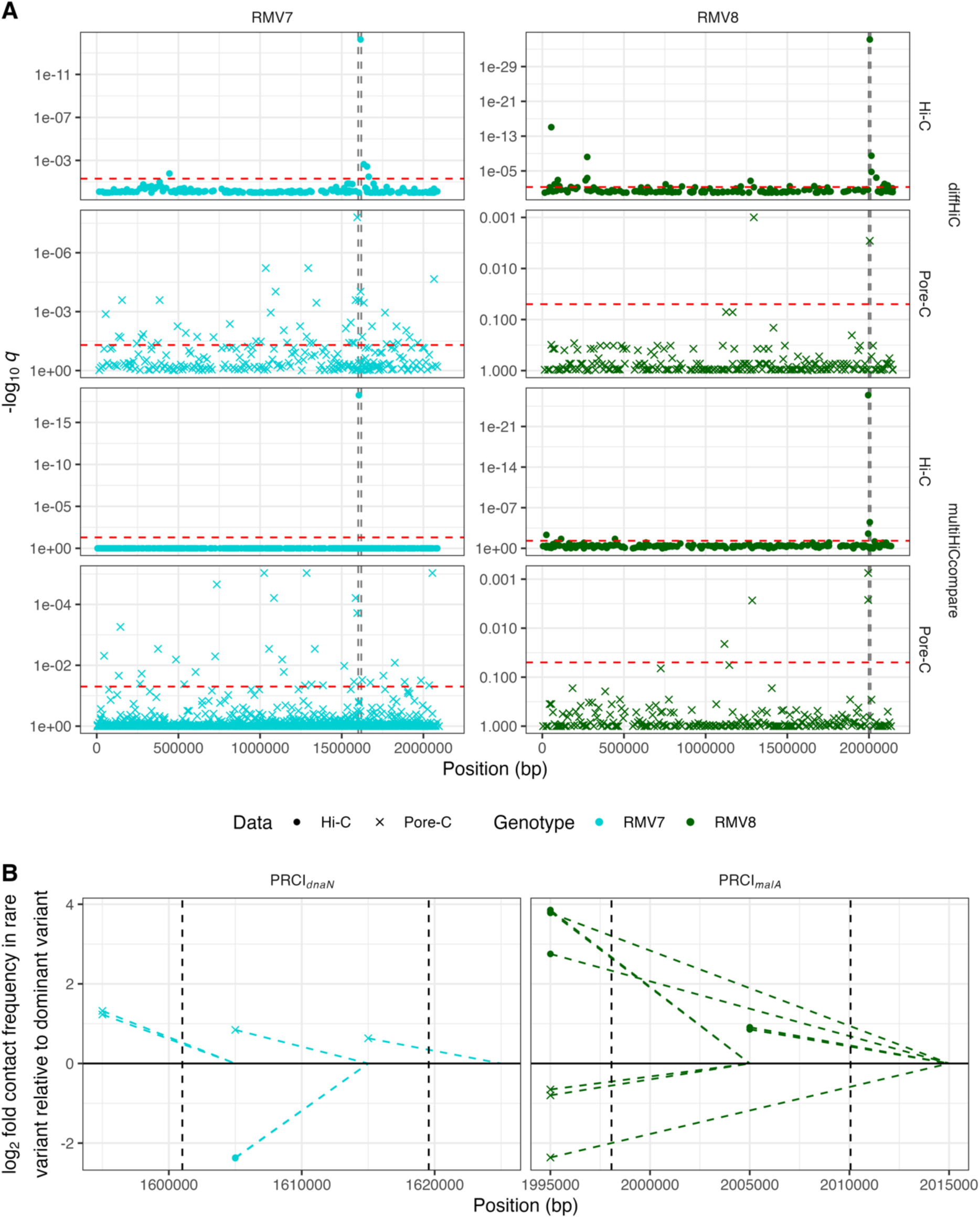
Genome-wide analysis of differential contact frequencies. **(A)** Manhattan plots comparing the contact densities between epigenetic variants. For each genotype, two different statistical methods (diffHiC and multiHiCCompare) were used to compare the three biological replicates generated using each technology (Hi-C and Pore-C) for both epigenetic variants. The negative logarithmic *q* values, corresponding to false discovery rates following a Benjamini-Hochberg correction for multiple testing, are shown for each locus across the genome. The threshold for genome-wide significance is shown by the red dashed horizontal line. The vertical black dashed lines show the boundaries of the variable PRCIs (PRCI*_dnaN_* in RMV7, and PRCI*_malA_* in RMV8) within the chromosomes. **(B)** Sites of differential contact frequencies identified by all four statistical comparisons within PRCI*_dnaN_* in RMV7, and PRCI*_malA_* in RMV8. Each point represents an interaction that was found to be significantly different between the corresponding dominant and rare variants following a correction for multiple testing. Its position on the horizontal axis represents the site of the interacting locus with the lower numerical position. The dashed line connects the point to the interacting locus with the higher numerical position. The vertical position of the point represents the base two logarithmic fold difference in the rare variant relative to the dominant variant, such that values greater than zero represent a higher contact frequency in the rare variant. The shapes of the points represent the method used to generate the data.

For RMV8, the only loci that were consistently identified as significantly differing between the two variants by all four comparisons were within, or adjacent to, PRCI*_malA_* [18]. The interacting loci, and the fold differences in contact frequency, were plotted against the annotation of this MGE (Fig. 2B). The hits were either between the flanking region and the element, or between the two flanks. However, while the spatial distribution and magnitude of the effects were similar across all comparisons, the Hi-C data suggested higher contact frequencies in RMV8_rare_, whereas the Pore-C data suggested higher contact frequencies in RMV8_domi_. To rule out sample misidentification as a cause of this discrepancy, it was verified that the patterns of contact frequencies at the *tvr* locus matched the arrangements characteristic of the variants, and that a simple analysis of contact frequency ratios within PRCI*_malA_* supported the genome-wide analysis results (Fig. S18). Detailed analysis of the Nanopore sequencing read lengths mapping to this region did not suggest this was an artefact of the Pore-C methodology preferentially sequencing shorter DNA fragments originating from the MGE (Fig. S19). Therefore the Hi-C and Pore-C analyses concurred that there was variation in the contact frequency within PRCI*_malA_*, but disagreed on the variant in which the contact density was highest.

The comparison of the RMV7 variants provided an opportunity to test whether this observation could be independently replicated, as the analyses of Hi-C data were consistent in identifying PRCI*_dnaN_* [12] as the region differing most significantly between RMV7_domi_ and RMV7_rare_ (Fig. 2A). This MGE was also the location of the most significant difference in contact frequencies between the variants in the diffHiC comparison of Pore-C datasets, in addition to being the site of multiple significant differences in the multiHiCcompare analysis of the same data (Fig. 1D). The detailed plot of these points relative to the PRCI*_dnaN_* annotation demonstrated these hits were either within the element, or between the element and its flanking regions (Fig. 2B). While the differences were again of similar magnitude and distribution across the two methods, the Hi-C and Pore-C data were also discordant in identifying the variant in which the contact frequency was highest. In this comparison, the Hi-C data suggested the contact densities were higher in RMV7_domi_, whereas the Pore-C data implied they were higher in RMV7_rare_. Therefore multiple significant differences in contact frequencies between both pairs of variants were within or flanking PRCIs, although the methods did not reach a consensus on the nature of the altered pattern of interactions.

### Changes in PRCI contact frequency do not reflect changes in XS

These discrepancies between the datasets for each variant could represent either a biological difference between the samples, or an artefact of the differing methodologies used to generate the sequence reads. Consequently, a generalised linear mixed-effects model was constructed that could resolve the effect of different genetic and biological variables. This model was fitted to the four combinations of sequencing methodology and genotype at a resolution of 10 kb using a zero-inflated Poisson distribution with a logarithmic link function (see Methods; Fig. S20-4). The fitted models were able to explain a substantial fraction of the variation in contact frequencies across loci and datasets (Fig. S25). The effect of distance between loci was estimated to be consistent between the genotypes, but contact frequencies declined more rapidly with distance in the Hi-C data, suggesting the Pore-C data were more efficient at detecting long-range interactions (Fig. 3A).

**Figure 3.**
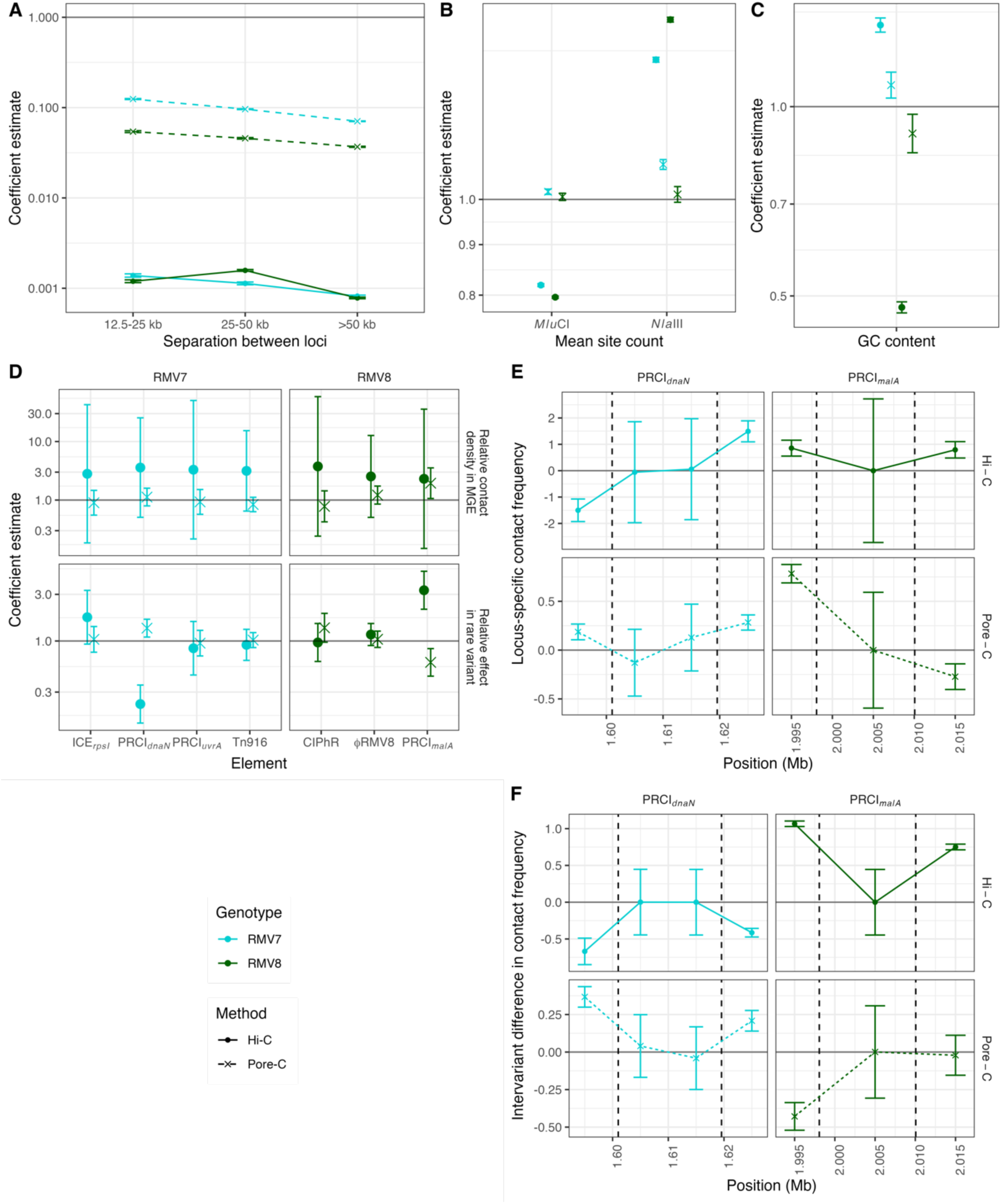
Statistical modelling of contact frequencies. A zero-inflated Poisson generalised linear mixed effects model was fitted to the four combinations of genotype and sequencing technology using contact matrices calculated at a resolution of 10 kb. The two genotypes are distinguished by colours, and the two technologies are distinguished by line and point styles. **(A)** Coefficients estimating the effect of separation between loci on contact frequencies, normalised to the expected contact frequency between neighbouring loci, shown by the horizontal line at a value of one. **(B)** Coefficients estimating the effect of *Mlu*CI and *Nla*III site densities on the detection of contacts; these coefficients reflect the impact of the number of sites per 1 kb. The horizontal line at one represents the coefficient value if the density of sites had no effect on the number of inferred contacts. **(C)** Coefficients estimating the effect of the fraction of bases (scaled upwards 10-fold to aid model fitting) that are guanine or cytosine (GC) on contact frequencies. **(D)** Coefficients estimating the contact frequencies within mobile genetic elements. The top panel shows the coefficients quantifying the relative contact frequencies in the MGEs, normalised to the frequencies observed across the rest of the chromosome, corresponding to the horizontal line at a value of one. The bottom panel shows the coefficients estimating the contact frequencies at the same loci in the rare variant relative to the dominant variant. **(E)** Locus-specific variation in the contact frequencies across the loci encompassing the two PRCIs highlighted in Fig. 2: PRCI*_dnaN_* within RMV7, and PRCI*_malA_* within RMV8. Data are plotted at 10 kb intervals. The black vertical dashed lines indicate the boundaries of the PRCIs. **(F)** Differences between the variants at the loci shown in (E).

To test whether any of the differential contact frequency results were artefacts of the restriction enzymes used to generate the samples, the model also estimated the sensitivity of the inferred contact frequencies to the density of sites targeted by these endonucleases (Fig. 3B). With the Pore-C data, there was no evidence of any strong relationship between restriction sites and inferred contact density. With Hi-C data, a higher density of *Nla*III sites was associated with an increased sensitivity for detecting contacts, which is consistent with the experimental protocol. However, there was an inverse relationship between the inferred contact density and the density of *MluC*I sites, despite this enzyme not being involved in sample preparation. Given the AT-rich nature of the *MluC*I sequence motif (AATT), such an observation may be attributable to Illumina sequencing’s bias towards DNA with a neutral GC content, which is not shared with Nanopore data [46]. However, contact frequencies did not exhibit an overall inverse relationship with GC content. Rather, contact frequencies increased with rising GC content in RMV7, but showed the opposite relationship in RMV8 (Fig. 3C). Both trends were more extreme for the Hi-C data than the Pore-C data. This suggested the Illumina data may be more affected by localised regions of extremely AT-rich sequence, correlating with the presence of *Mlu*CI sites, rather than the mean GC content averaged over a 10 kb locus. Nevertheless, no consistent evidence could be found of a pneumococcal XS system that preferentially condensed AT-rich sequences [47].

A more specific test for XS was quantifying the contact density across the MGEs within each of the genomes, relative to the rest of the chromosome. For RMV7, neither the Hi-C or Pore-C data found any evidence of elevated contact densities across the identified MGEs, with the confidence intervals of the level of contact density relative to the rest of the genome always spanning one (Fig. 3D). When comparing the variants, differences in the density of contacts were only inferred for PRCI*_dnaN_*. In further agreement with the differential contact analyses (Fig. 2B), the contact frequencies were higher for PRCI*_dnaN_* in RMV7_rare_ for the Pore-C data, but lower in the Hi-C data. For RMV8, the Illumina Hi-C data again found no evidence of the ϕRMV8 or PRCI*_malA_* MGEs being associated with elevated contact densities, although there was some evidence of higher contact frequencies within the latter in the Pore-C data. However, the Pore-C data also suggested that contact frequencies within PRCI*_malA_* were reduced in the rare variant, suggesting there was no consistent evidence for increased contact densities across both variants. By contrast, the Illumina Hi-C data suggested that PRCI*_malA_* was associated with higher contact densities in RMV8_rare_. Therefore the model outputs were consistent with the differential contact analysis in both genotypes, and there was no evidence of elevated contact densities within the pneumococcal MGEs that could provide evidence of XS.

To test whether there was evidence of XS at any location within MGEs, locus-specific effects on contact frequencies were estimated for the sites within and flanking PRCI*_dnaN_* and PRCI*_malA_* (Fig. 3E). There was no evidence of heterogeneity within the elements, although the size of PRCI*_malA_* meant it contained only a single locus, making the elevated contact density inferred for this element in the Pore-C data highly susceptible to confounding by locus-level effects (Fig. 3D). There was also little variation between the variants at these sites (Fig. 3F), which is consistent with the differences being captured by the MGE-level variables (FIg. 3D). However, variation was evident in the flanking regions, the direction of which mirrored the alterations in the elements themselves (Fig. 3D). Therefore the differential contact densities at the PRCIs was not caused by changing levels of XS-driven condensation, but instead reflected altered interactions between the MGEs and the proximal regions of the host chromosome.

### Contact frequency changes are associated with altered PRCI activity

Such an observation suggested the differences between the variants may reflect changes in excision, replication and activation of the PRCIs. Given the conflicting data between the Hi-C and Pore-C methods, previously-collected RNA-seq data were used to identify the variants in which the MGEs were more active. PRCI*_dnaN_* had previously been identified as being transcribed at a significantly higher level in RMV7_domi_ than RMV7_rare_ during early exponential phase using both RNA-seq (Fig. 4A) and quantitative reverse transcriptase PCR (qRT-PCR) data [12]. Analogously, the PRCI*_malA_* genes were generally more highly expressed in RMV8_rare_ than RMV8_domi_ (Fig. 4B), although these differences were not consistent enough to reach genome-wide significance [18]. However, no difference was observed when the *tvr* loci of the variants were removed (Fig. 4C), indicating the differences might be associated with epigenetic modifications [18].

**Figure 4.**
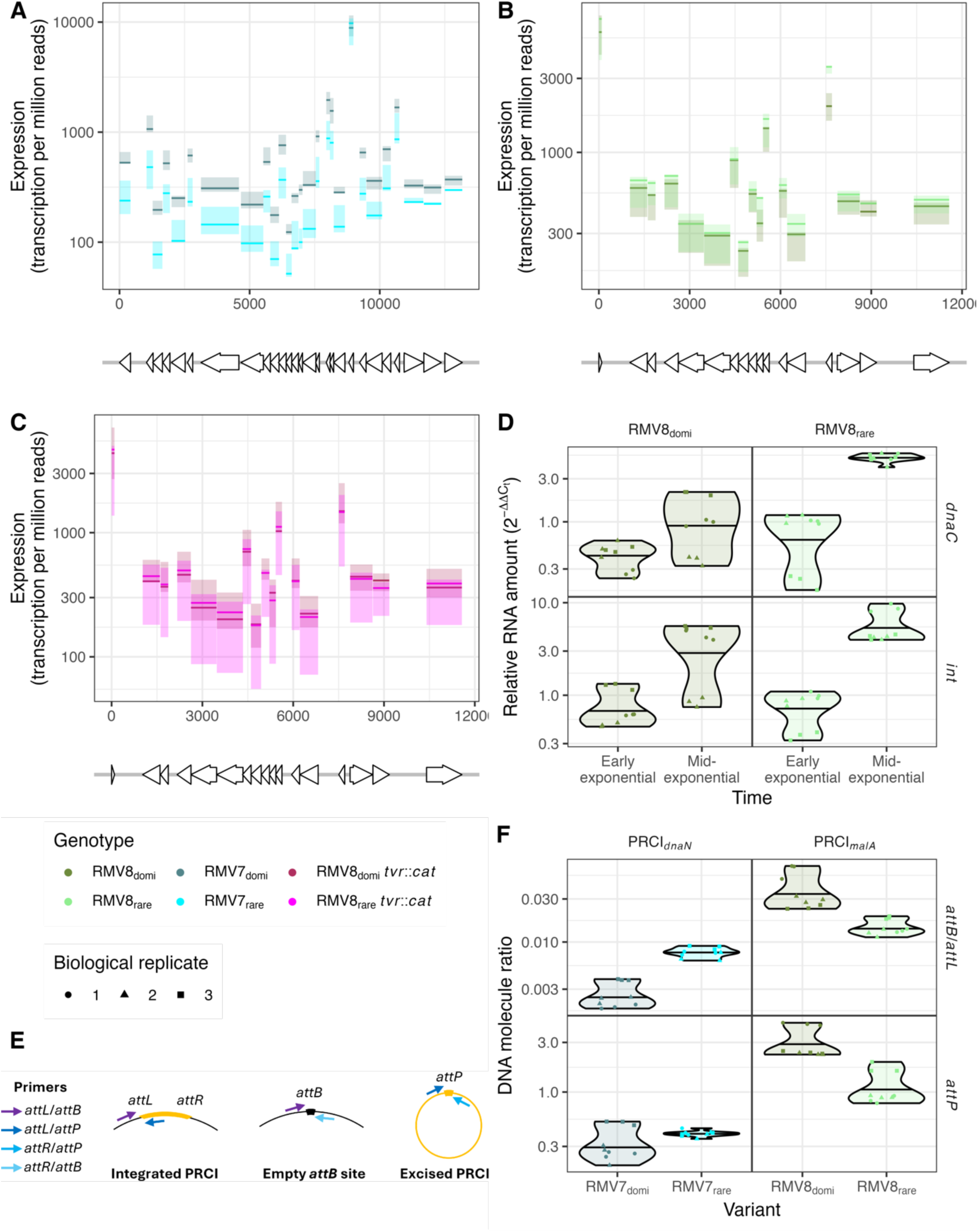
**Differential activity of PRCIs between variants**. **(A)** Quantification of PRCI*_dnaN_* expression in RMV7_domi_ and RMV7_rare_ using previously-published RNA-seq data [12]. The annotated protein coding sequences (CDSs) within the PRCIs are shown by the arrows at the bottom of the panel, with their orientation indicating the direction in which they are transcribed. The lines in the main panel show the level of expression estimated for each CDS for each of the three independent biological replicates as normalised transcripts per million reads. The solid line indicates the median value for each variant, and the shaded area indicates the range between the maximum and minimum values. **(B)** Quantification of PRCI*_malA_* expression in RMV7_domi_ and RMV7_rare_ using previously-published RNA-seq data [18], as displayed in panel (A). **(C)** Quantification of PRCI*_malA_* expression in RMV7_domi_ *tvr*::*cat* and RMV7_rare_ *tvr*::*cat* using previously-published RNA-seq data [18], as displayed in panel (A). These two mutants do not have functional *Spn*IV RMSs. **(D)** Quantification of PRCI*_malA_* expression in RMV7_domi_ and RMV7_rare_ using qRT-PCR assays of transcription of an integrase (*int*) and replication (*dnaC*) CDS. Three technical replicates are shown for each of three biological replicates, as indicated by the shapes of the points. The violin plot summarises their distribution, with the median indicated by the horizontal line. Measurements were made at both early and mid-exponential timepoints. **(E)** Diagram showing the primers used to quantify the integration, excision and replication of the PRCIs. **(F)** Differences in integration, excision and replication of the PRCIs between variants. Data are shown as in panel (D). For both PRCI*_dnaN_*, in RMV7, and PRCI*_malA_*, in RMV8, the level of excision was quantified as the ratio of *attB* sites to *attL* sites A higher ratio indicates a greater level of excision. Similarly, the levels of circularised PRCIs were quantified by measuring the concentration of *attP*.

To validate whether transcription of PRCI*_malA_* substantially differed between the variants, qRT-PCR was used to assay the expression of *int*, encoding the element’s integrase, and *dnaC*, encoding a DNA replication protein, during early exponential (OD_600_ = 0.2) and mid-exponential (OD_600_ = 0.5) phase growth (Fig. 4D). Although no substantial difference was evident in early exponential phase, the transcription of the replication gene rose to be ∼4-fold higher in RMV8_rare_ than RMV8_domi_ during mid-exponential phase, consistent with the higher expression of these genes in the same variant in the RNA-seq data (Fig. 4B). This suggested that PRCI*_dnaN_* and PRCI*_malA_* were less frequently transcribed in RMV7_rare_ and RMV8_domi_, respectively. Hence transcription, as well as contact densities, differed between epigenetic variants at these PRCIs.

The readily detectable levels of gene expression, which changed in response to growth, were also consistent with an absence of XS. Instead, these transcriptional signals indicated that the PRCIs were likely to be excising and replicating, which would explain the differences in contacts within these elements, and between them and their flanking regions [48]. Therefore qPCR assays were used to measure the ratios of excised to integrated elements, and the copy number of circularised elements within the cell (Fig. 4E,F). In RMV7, the ratio of excised to integrated PRCI*_dnaN_* was significantly higher in RMV7_rare_, although there was no substantial difference in the number of circularized elements within the cell. In RMV8, both the ratio of excised to integrated elements, and the frequency of circularised PRCI*_malA_* genomes, were significantly higher in RMV8_domi_. Therefore the altered contact frequency patterns between variants likely represented changes in the topological configuration of the elements.

For such a topological change to fully explain the significant differences of opposing magnitude in the Hi-C and Pore-C comparisons, it would need to be possible for differences in PRCI circularization to be metastable [49]. Under such circumstances, a variant may exhibit levels of PRCI circularization and replication that are stable over enough bacterial generations such that separate biological cultures grown in parallel have consistent properties. Yet the conflicting results between analyses demands that it is also possible that re-isolation of the variant produces a genotype with a different level of PRCI activity that is similarly persistent.

To test this, three independent colonies were cultured for each variant of RMV7 and RMV8. Each of these cultures was frozen, used to inoculate overnight growth, then diluted into three parallel tubes of fresh media until they reached exponential phase growth. Measurements of the level of excision, and extracellular circularization, were generally consistent across the parallel triplicate cultures inoculated from the same isolate (Fig. 5A). However, there was substantial variation between cultures derived from different colonies for RMV8. These results also contrasted with the previous qPCR assays, which were performed on the same isolate used for the Pore-C analysis (Fig. 4F). For RMV7, the results were more consistent across the colonies representing each variant (Fig. 5A). However, RMV7_domi_ was generally associated with the higher median copy number across the replicates in this experiment, whereas this value had been higher for RMV7_rare_ in the previous assay (Fig. 4F). Therefore PRCI copy number levels varied stochastically between even isogenic isolates, but are metastable during culturing in liquid media.

**Figure 5.**
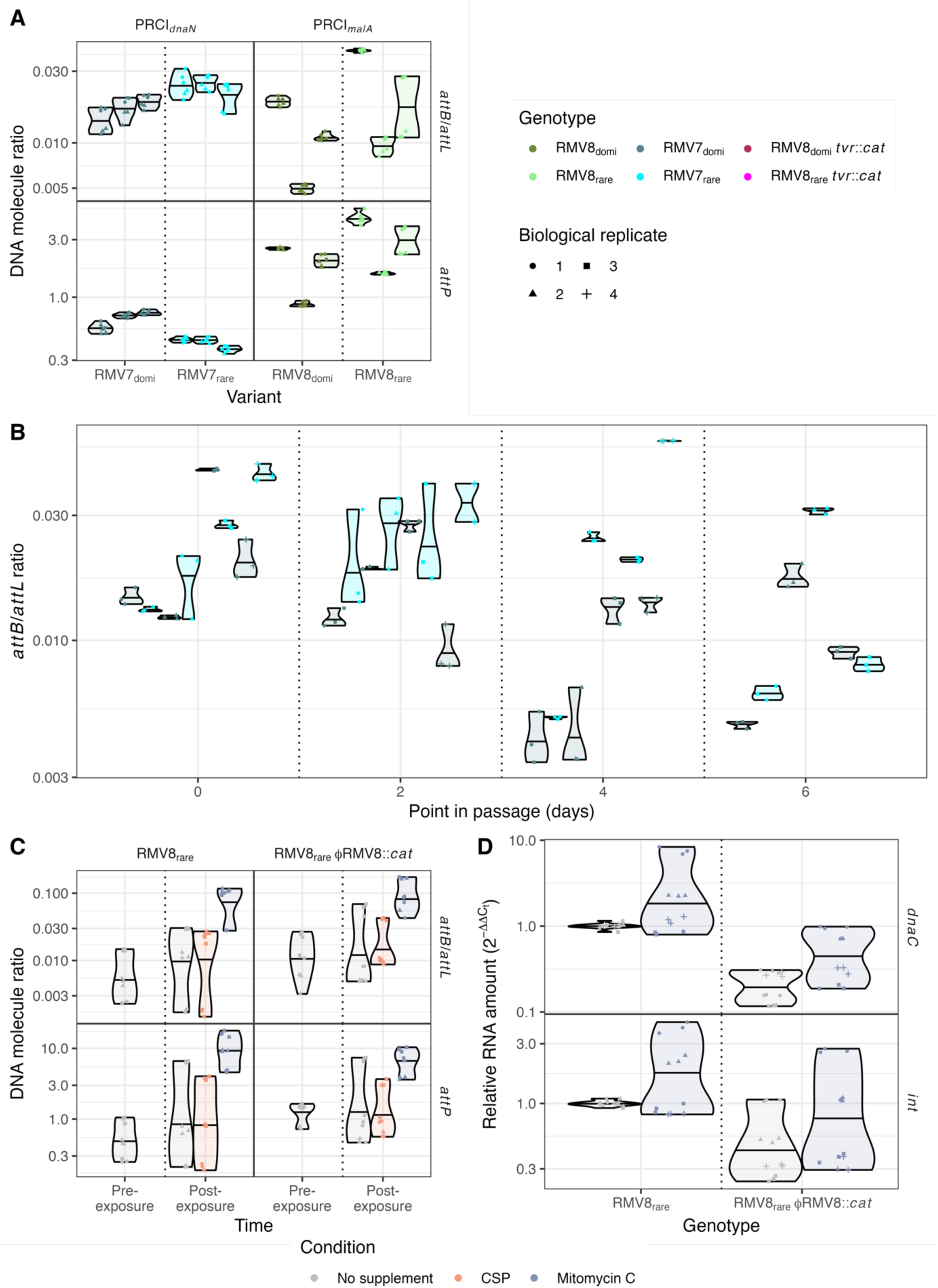
**Metastability and inducibility of PRCI activity**. **(A)** Violin plot showing the variation in PRCI topology between cultures inoculated with independently-isolated colonies of the four variants. Three biological replicate cultures were grown from each of three independently-isolates colonies for each variant. Each violin plot represents a single colony, and describes the distribution of three technical replicate measurements for an individual biological replicate, with the shape of each point indicating the biological replicate to which they relate. The horizontal line across each violin represents the median value. **(B)** Violin plot showing the variation in PRCI*_dnaN_* excision in a six-day passage of RMV7 genotypes. Four independent mutants, in which the RMV8_domi_ or RMV8_rare_ *tvr* locus had been introduced into a *tvr*-null RMV8 genotype, were passaged in parallel [12]. DNA was sampled from a single biological replicate of each mutant at the start of the passage, and after two, four and six days. Each violin summarises three technical replicate measurements of a single biological replicate. **(C)** Violin plot showing the effect of inducing stimuli on the topology of PRCI*_malA_*. The excision and replication of PRCI*_malA_* following exposure to competence stimulating peptide and mitomycin C were assayed two hours after exposure in early exponential growth. These experiments used both RMV8_rare_, and a mutant derivative lacking the ϕRMV8 prophage. Data are shown as in panel (A). **(D)** Violin plot quantifying the effect of mitomycin C on PRCI gene expression. The expression of the *int* and *dnaC* genes of PRCI*_dnaN_* were assayed after two hours of exposure to mitomycin C, in both RMV8_rare_ and a mutant derivative lacking the ϕRMV8 prophage. Data are shown as in Fig. 4.

### PRCI excision and expression are separately regulated

The variation in contact frequency and copy number between isolates of the same variant contrasted with the consistent differences in PRCI expression between RNA-seq and qRT-PCR data experiments (Fig. 4B,D) [12]. Correspondingly, previous passages of RMV7 genotypes, in which *tvr* loci with differing arrangements were independently introduced, found phenotypic and expression differences were stable over multiple days [12]. The topological arrangement of PRCI*_dnaN_* was assayed in the same samples (Fig. 5B). This demonstrated substantial variation in the integration-excision dynamics of this element across isolates of both variants. As these samples were directly transferred between liquid cultures, this demonstrated the changes in PRCI excision did not depend on culturing on solid media. Hence differences in PRCI expression may be stable, despite stochastic variation in the topology of the element.

This implied the excision and transcription of PRCIs may be differentially regulated, consistent with PRCI gene expression in at least some samples being highest in the variants in which these elements were most frequently integrated into the chromosome (Fig. 4D,F). This was tested using RMV8, which carries both the ɸRMV8 prophage and PRCI*_malA_* [18]. RMV8_rare_ cells were exposed to competence stimulating peptide (CSP) and mitomycin C (MMC), both of which activate the ɸRMV8 prophage [20]. These assays were conducted with RMV8_rare_ and a mutant in which ɸRMV8 had been replaced with a chloramphenicol acetyltransferase (*cat*) resistance marker, to ascertain whether the PRCI’s activity was affected by the prophage (Fig. 5C). Both the level of excision, and the concentration of circularised PRCI elements, rose in response to MMC, but not CSP. This behaviour was independent of the presence of a prophage within the same cell. This suggests the excision of PRCI*_malA_* is controlled by the MGE itself in response to similar stimuli that activate prophage.

A qRT-PCR assay was then used to test whether MMC also activated PRCI*_malA_ int* and *dnaC* gene expression in both RMV8_rare_ and RMV8_rare_ ɸRMV8::*cat* (Fig. 5D). An inconsistent rise in the transcription of both genes was recorded in MMC supplemented cultures, correlating with the elevated copy number of the element (Fig. 5C). However, independently of treatment with MMC, transcription levels were consistently lower in the absence of the prophage (Fig. 5D). This effect was greatest for the *dnaC* replication gene, the median expression of which was 4.8-fold lower in the absence of ɸRMV8 in unsupplemented media, and 3.4-fold lower in the absence of ɸRMV8 in MMC-supplemented cultures. This is consistent with the PRCI’s dependence on the intact prophage for its ability to transmit between host cells [50]. Therefore expression of the PRCI is regulated by additional factors, associated with the presence of prophage, that do not affect excision.

### PRCIs do not cause a substantial fitness cost to their host cells

To test whether this regulation of PRCI activity limited their cost to their host cell, the growth of mutants in which PRCIs had been replaced with a Janus cassette [51] was compared to the corresponding unmodified genotypes in unsupplemented media (Fig. 6A). Neither RMV7_rare_ PRCI*_dnaN_*::Janus, nor RMV8_rare_ PRCI*_malA_*::Janus, exhibited detectable changes in growth in unsupplemented media relative to the variant from which they were derived. By contrast, removal of the prophage ϕRMV8 substantially increased the growth of RMV8_rare_ (Fig. 6B), consistent with the behaviour of a similar RMV8 prophage null mutant [20]. The deletion of PRCI*_malA_* from this non-lysogenic mutant caused no additional change. Therefore PRCIs cause a substantially smaller impairment of host pneumococcus growth than prophage in the absence of an activating stimulus.

**Figure 6.**
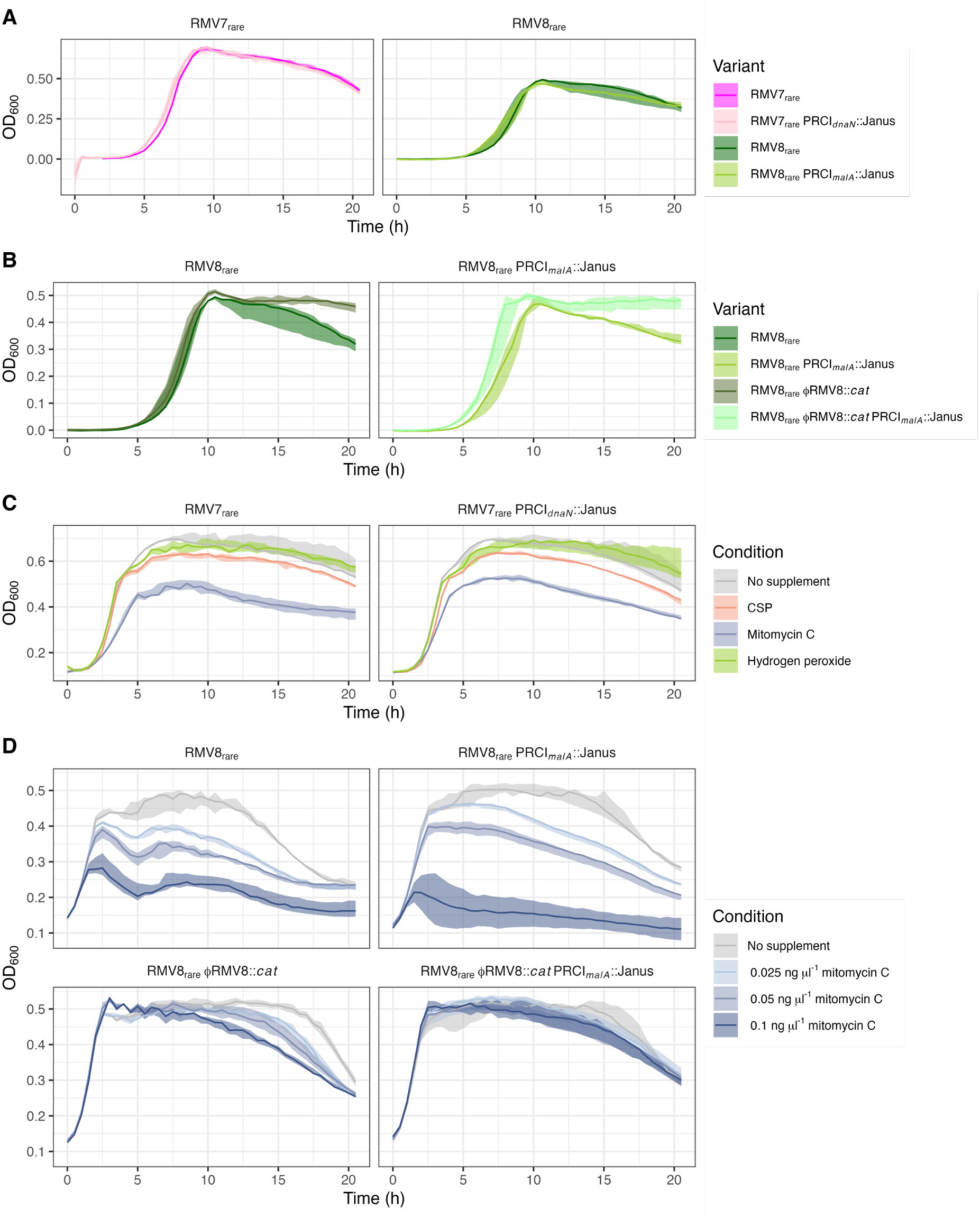
The impact of induced and uninduced MGEs on the growth of RMV7 and RMV8 genotypes. Each panel shows the 20 h growth curve of a set of pneumococcal genotypes, measured through assaying the optical density (OD_600_) at 30 min intervals. In each plot, the solid line shows the median, and the shaded region shows the full range observed across three biological replicates. **(A)** A comparison of the effect of removing PRCI*_dnaN_* from RMV7_rare_ and removing PRCI*_malA_* from RMV8_rare_. **(B)** A comparison of the effect of removing ϕRMV8 from RMV8_rare_ and RMV8_rare_ PRCI*_dnaN_*::Janus. **(C)** A comparison of the effect of exogenous stimuli on the growth of RMV7_rare_ and RMV7_rare_ PRCI*_dnaN_*::Janus. **(D)** The effect of increasing concentrations of mitomycin C on RMV8_rare_ and mutant derivatives lacking ϕRMV8 or PRCI*_malA_*.

Growth assays were then used to test whether this changed when PRCIs were induced to excise from the chromosome by MMC during early exponential growth. Both RMV7_rare_ and the corresponding PRCI*_dnaN_*::Janus grew similarly in the presence of MMC, consistent with the MGE not reducing host cell fitness upon excision (Fig. 6C). Although MMC did cause a dose-dependent reduction in the density of RMV8_rare_ cultures, this sensitivity was substantially reduced in a non-lysogenic ϕRMV8::*cat* mutant (Fig. 6D). By contrast, removal of PRCI*_malA_* did not affect MMC sensitivity in either the wild type or non-lysogenic RMV8 cells. Therefore stimulating the excision and circularisation of PRCIs neither adversely affect pneumococcal host cell growth, nor interfered with the lysis of cells following prophage induction.

## Discussion

This application of both Hi-C and Pore-C methods to epigenetic variants of two *S. pneumoniae* genotypes provides an overview of the pneumococcal chromosome’s organisation. Both methods concurred that the chromosomal contacts were dominated by short-range interactions, consistent with a simple, locally-condensed structure. However, the resolution at which these could be studied was limited by the density of digestion and proximity re-ligation events. Future studies, using refined sample preparation protocols, may enable higher-resolution analyses of bacterial chromosomes.

At larger scales, some similarities were observed with data from *B. subtilis*, another species within the Bacillota phylum [52], and a recently-published study of *S. pneumoniae* R6 [53]. Both methods found evidence of an origin domain, indicative of this region having a globular conformation, which included constrained contact patterns (Fig. 1D) that likely reflected the positioning of *parS* sites, as seen in *B. subtilis* [52] and *S. pneumoniae* [53]. The Pore-C data also identified a secondary diagonal signal of long-range interactions that was orthogonal to that formed by the contacts between neighbouring loci, previously determined as being indicative of a close association of the chromosome arms (Fig. 1C) [52]. However, there was no evidence of the chromosome interaction domains that have been noted not only in *B. subtilis* [52], but also *E. coli* [54,55], *Caulobacter crescentus* [56] and *Mycoplasma pneumoniae* [57]. This absence of discrete domains concurs with the recent study of *S. pneumoniae* R6 [53]. It may be that the relatively small pneumococcal chromosome, with strong coding biases on each replichore [58], is capable of orchestrating concurrent transcription and regulation without substantial higher-order chromosomal folding.

This comparatively simple structure may reflect the apparent paucity of NAPs in pneumococci relative to *E. coli* and *B. subtilis*, with only the histone-like protein HU (or HlpA) characterised in *S. pneumoniae* [59–62], and the relatively small number of MGEs in a typically pneumococcal genome [4]. The lack of characterised orthologues of many of the proteins involved in XS is consistent with the absence of condensed regions that form boundaries between large domains in other species [52], and the extensive disruption to pneumococcal chromosome conformation associated with the depletion of HlpA [53]. Nevertheless, the epigenetic variants of RMV7 and RMV8 were each distinguished by variable contact density at the sites of PRCI integration (Fig. 2A). However, the approximately two-fold increases in contact density at these elements (Fig. 2B) was consistent with the *attP* locus of the circularised form having a similar copy number to the chromosome itself (Fig. 4E, Fig. 5A), effectively doubling the presence of the PRCI sequences. Hence the alternative explanation of condensation by XS was unlikely. While it is possible that the high density of short-range contacts identified chromosome-wide in our analyses prevents the identification of greater condensation of loci, the absence of XS also explains the absence of any detectable increase in contact density in AT-rich regions, or within other MGEs (Fig. 3C, D).

Furthermore, a lack of XS is consistent with the continued expression, excision and activation of these PRCIs (Fig. 4, 5), and the larger ϕRMV8 prophage (Fig. 6), even in the absence of inducing stimuli [18,20]. Such a difference in genome biology from *B. subtilis* and *E. coli* suggests a fundamentally distinct interaction between MGEs and host cells in these species. One possible explanation is that XS proteins have an important role in controlling the activity of plasmids [63]. Such elements are common in *E. coli* [64], found in ∼10% of *B. subtilis* isolates [65], but rarer in *S. pneumoniae* [4]. Hence pneumococci may have adopted a different strategy for reducing the fitness cost or transmissibility of their primarily chromosomally-integrated MGEs.

That the only consistent significant difference in contact frequencies between the variants of RMV8 was caused by PRCI*_malA_* (Fig. 2A) contrasted with the previous RNA-seq analysis, which only identified *comC* and ϕRMV8 as sites of significantly-differing transcriptional variation between RMV8_domi_ and RMV8_rare_ [18]. However, the excision of ϕRMV8 is linked to its increased expression, typically resulting in cell lysis [20]. By contrast the excision of PRCI*_malA_* is reversible, without substantial fitness cost to the host (Fig. 6), meaning these episomes are always likely to be hotspots of variation in copy number even in cells growing in the same culture, thereby resulting in differential contact densities. Additional significant differences in contact density were also identified across the genome in the Pore-C data, with greater variation in RMV7 than RMV8. This is consistent with the more widespread differences in transcription between RMV7_domi_ and RMV7_rare_ than between the RMV8 variants [12,18], although it may also reflect the higher and more even coverage across the RMV7 Pore-C datasets (Fig. S8) providing greater power for detecting such variation. Hence alterations in chromosomal conformation remains a feasible mechanism to explain how distinct patterns of epigenetic DNA modification can affect transcription. However, this observed correlation cannot establish causation without higher-resolution 3C data, and further experimental work that can change both the DNA methylation patterns, and the organisation of the genome.

However, the DNA modifications could not account for the differences in excision dynamics, which were variable between different isolates of the same variant (Fig. 5A). This diversity likely represents the PRCIs engaging in “bet hedging” behaviour [12,66], as integrated elements are stably inherited following chromosomal replication, but are at risk of deletion by homologous recombination [67]. Hence the vertical inheritance of PRCIs is likely to be maximised by existing as both chromosomally-integrated and episomal forms. That the *attB*/*attL* ratios were below one suggested most *attB* sites contained a chromosomally-integrated form (Fig. 4D), ensuring they will be passed to daughter cells in clonally-replicating populations. However, the *attP* levels were in the range 0.3-5, implying that circular forms were also widespread, meaning the PRCI can surivive even if the chromosomally-integrated copy is deleted by recombination. Therefore these episomal dynamics can help explain the stable inheritance of PRCIs by strains over decades, in contrast to the transient associations between prophage and host genotypes [4].

This accounts for why it is adaptive for the PRCIs to increase their excision in response to the detection of RecA-coated nucleoprotein filaments (Fig. 5C), which are generated by MMC, and mediate homologous recombination [20]. Yet PRCI excision does not cause the same fitness cost to the cell as prophage activation, because it cannot lyse the host in the absence of an intact “helper” phage (Fig. 5D). Therefore it is also adaptive for expression to be regulated in response to prophage activity, as was apparent in RMV8. This observed interdependence between the MGEs explains the rise in PRCI*_malA_* transcription in mid-exponential phase (Fig. 4F), as this is when ϕRMV8 is most active [68]. This distinction between the regulation of excision and activation means the former can occur stochastically, even in the absence of a prophage, as in RMV7. The observed epigenetic inheritance of a variable state suggests that the levels of excision are determined by the concentration of regulatory molecules being passed on from mother to daughter cells [69], although no mechanistic details were analysed in this study. Hence the discrepancy between the PRCI copy numbers detected in the Hi-C and Pore-C data is indicative of these genomes being dynamic systems, the behaviour of which is determined by metastable regulatory switches, which cannot by fully described by a single sequence alone.

Hence Pore-C will reveal additional heterogeneity within species, strains, and even clonal bacterial populations, with identical genomes being differentiated by their topology and copy number of individual elements. Although our analysis used data from two flow cells, one was sufficient to identify the key differences between the studied genotypes, meaning this method can be implemented by any laboratory with the capacity to use sequencing platforms on the scale of a MinION. Hence future applications of this method to species with a greater diversity of replicons, NAPs and XS systems promises to aid our understanding of the diversity of bacterial genome biology across species.

## Methods

### Culturing of *S. pneumoniae*

All *S. pneumoniae* strains (Table S1) were cultured at 35°C in a 5% CO_2_ atmosphere. Culturing on solid media used Todd-Hewitt broth (Sigma-Aldrich) supplemented with 0.5% yeast extract (Sigma-Aldrich), 1.5% agar (Sigma-Aldrich) and 200 U ml^−1^ catalase (Sigma-Aldrich). To select desired mutant genotypes, media were additionally supplemented with antibiotics: streptomycin (Sigma-Aldrich) or kanamycin (Sigma-Aldrich) at 400 µg ml^-1^, or chloramphenicol (Sigma-Aldrich) at 4 µg ml^-1^.

Unless stated otherwise, liquid culturing utilised a ‘mixed’ media consisting of Todd-Hewitt broth supplemented with 0.5% yeast extract and Brain Heart Infusion broth (Sigma-Aldrich) mixed at a 2:3 ratio [12]. For comparing PRCI dynamics between variants, samples were collected during mid-exponential growth, at an optical density at 600 nm (OD_600_) of ∼0.6. For measuring the PRCI responses to stimuli, either unsupplemented media, MMC (850 ng), or competence stimulating peptide (CSP; 8.75 µg) was added to 7 ml of bacteria grown to an OD_600_ of ∼0.2. The added CSP was appropriate for inducing competence in each genotype: CSP1 for RMV7, and CSP2 for RMV8. The cultures were then incubated at 35°C with 5% CO_2_ for 2 h. Subsequently, 5 ml of culture was used for RNA extraction, and 2 ml was centrifuged at 3220 *g* for 10 min to produce a cell pellet for DNA extraction.

To measure growth curves, 2 x 10^4^ cells from titrated frozen glycerol stocks were grown in mixed liquid media in 96-well microtiter plates at 35°C with 5% CO_2_ for 20 h. For all experiments measuring bacterial sensitivity to inducing stimuli, 200 µl of bacterial cultures with an OD_600_ between 0.15 and 0.25 were mixed with MMC (10 ng), CSP (75 ng), or H_2_O_2_ (2 ng), then grown in microtiter plates at 35°C with 5% CO_2_ for 20 h. The OD_600_ was measured at 30 min intervals using a FLUOstar Omega microplate reader (BMG LABTECH). Three or four replicate wells were assayed for each tested genotype and condition in each experiment.

### DNA purification and PCR amplification

Following overnight incubation, bacteria were pelleted by centrifugation at 3220 *g* for 10 min. Supernatants were discarded, and bacterial pellets were then resuspended in 480 µl of lysis buffer (Promega) and 120 µl of 30 mg ml^-1^ lysozyme (Sigma-Aldrich). Samples were incubated at 35°C for 45 min and centrifuged for 2 min at 8000 *g*. Bacterial genomic DNA was then extracted using Wizard Genomic DNA Purification Kit (Promega), following the manufacturer’s instructions. For experiments requiring subsequent PCR product purification, 1000 ng of genomic DNA, 25 µl of 2x DreamTaq Master Mix (Thermo Fisher), nuclease-free water (Thermo Fisher) and 2 µl each of the 10 µM forward and reverse primers listed in Table S4 were used for PCR amplification in a total reaction volume of 50 µl. Otherwise, 500 ng of genomic DNA, 7.5 µl of 2x DreamTaq Master Mix (Thermo Fisher), 1 µl each of a 10 µM forward and reverse primers (Table S2), and nuclease-free water were combined in a total reaction volume of 15 µl.

### Construction of the *S. pneumoniae* RMV8 PRCI*_malA_* mutant

To generate the PRCI*_malA_*::Janus construct, the ∼1 kb region flanking upstream (UP) and downstream (DOWN) of PRCI*_malA_* were amplified by PCR using primers that inserted restriction enzyme sites on the internal sides of each region. The UP region was generated with primers malA_KO_UP_For and malA_KO_UP_Apal, whereas the DOWN region was generated with primers malA_KO_DOWN_For_BamHI and malA_KO_DOWN_Rev. The Janus cassette was amplified with the corresponding primers with added restriction sites (Table S4). Purified PCR amplicons were then obtained via gel electrophoresis with a 1% agarose gel dyed with SYBR Safe (Thermo Fisher) in Tris/Borate/EDTA buffer, followed by gel extraction with a GenElute Gel Extraction Kit (Sigma-Aldrich).

Following extraction, purified PCR amplicons were digested at 37°C for 2 h with the appropriate restriction enzyme: ApaI (Promega) for UP, and BamHI (Promega) for DOWN. Digested UP and DOWN products were then mixed with the digested Janus cassette at a 2:2:1 ratio and incubated with T4 DNA ligase (New England Biolabs) at room temperature (RT) overnight. The ligation mix was then used as the template for a touchdown PCR, and the amplified PRCI*_malA_*::Janus construct was purified by separation through electrophoresis on a 1% agarose gel, and extraction using a QIAquick Gel & PCR Cleanup kit (Qiagen). To transform recipient bacteria (RMV8_rare_ or RMV8_rare_ ϕRMV8*::cat*) with the PRCI*_malA_*::Janus construct, a 1 ml sample of a bacterial culture at an OD_600_ of ∼0.2 was incubated with 5 µl of 500 mM CaCl_2_ (Sigma-Aldrich), 1250 ng of CSP2, and 1 µg of the PRCI*_malA_*::Janus construct at 35°C for 2 h. Samples were then spread onto solid media supplemented with antibiotics (400 µg ml^-1^ of either kanamycin or streptomycin) to select for resistant transformants that grew after incubation at 35°C with 5% CO_2_ for at least 16 h.

### RNA purification and reverse transcription

A 5 ml sample of a bacterial culture was mixed with an equal volume of RNAprotect (Qiagen) and incubated at room temperature for 5 min. RNA samples were extracted using the SV Total Isolation System (Promega) according to the manufacturer’s instructions, with the exception that the initial extraction of RNA required treating cell cultures with 480 μL 30 mg mL^−1^ lysozyme (Promega) for 30 min at 35 °C. The purified RNA was treated with amplification-grade DNase I (Invitrogen), then used in reverse transcription reactions using the First-Strand III cDNA synthesis kit (Invitrogen). Each reaction used 0.2 μg RNA, 100 units of the SuperScript III reverse transcriptase, 1 μL of 100 μM random hexamer primers (Thermo Fisher Scientific) and 1 μL of 10 mM dNTP mix (Bioline). After a 5 min annealing period at 25 °C, reverse transcription was conducted at 50 °C for 30 min, then 55°C for 30 min. The reverse transcriptase was then heat inactivated at 70 °C for 15 min.

### Quantitative PCR

The oligonucleotides used for qPCR were designed to generate amplicons that were 150-200 bp in length, and are listed in Table S4. Template cDNA was diluted in a 1:25 ratio with nuclease-free water (Qiagen). Each 15 μL reaction used 3.75 μL of DNA template (either genomic DNA or cDNA), 0.75 μL of 10mM forward and reverse primer solutions (Invitrogen), 7.5 μL of PowerUp SYBR Green Master Mix (Thermo Fisher Scientific) and 2.25 μL of DNase-free and RNase-free water. Reactions were run using MicroAmp Optical 96-well reaction plates (Thermo Fisher Scientific) and the QuantStudio 7 Flex Real-Time PCR System (Applied Biosystems). The mixtures were initially heated to 50 °C for 2 min. Subsequently, 40 amplification cycles were run, each consisting of denaturation at 95 °C for 15 s, followed by annealing and elongation at 60 °C for 1 min. The purity of the amplicon at the end of the reaction was assessed through a melt curve analysis. In all experiments, three technical replicate measurements were made for each of three biological replicates.

The expression of PRCI*_malA_* genes was quantified using the ΔΔCt method. The *rpoA* gene was used as the reference gene, against which target gene expression levels were normalised to yield a ΔCt value [12]. The ΔCt values were calculated as the difference between the mean Ct of the target gene and the reference gene across technical replicates within the same biological replicate. The ΔΔCt values were then calculated relative to the reference ΔCt, which was that of RMV8_rare_ at an OD_600_ of 0.2. The fold gene expression difference between genotypes was then quantified as 2^-ΔΔCt^.

For the quantification of PRCI dynamics, the amplicons produced across the integration site, and the *rpoA* reference gene, were generated from genomic DNA (Table S2). The concentration of each purified PCR product was measured using a Qubit Broad Range Kit and Qubit 4 Fluorometer (Thermo Fisher). The Ct values for copy numbers of 3×10^3^, 3×10^4^, 3×10^5^ and 3×10^6^ of these products were used to generate a standard curve to convert the Ct values from experiments into absolute copy numbers of DNA molecules.

### Construction of Hi-C and Pore-C sequencing libraries

Three independent cultures of each of the four variants (*S. pneumoniae* RMV7_domi_, RMV7_rare_, RMV8_domi_ and RMV8_rare_) were grown to mid-exponential phase, corresponding to an OD_600_ of ∼0.60. The samples processed for Illumina Hi-C and Pore-C analyses were cultured independently. To crosslink DNA and proteins, cells were mixed with 1% v/v formaldehyde (Sigma-Aldrich) for 30 min at RT, followed by a 30 min incubation at 4°C. The formaldehyde was then quenched through the addition of 1 ml 2.5 M glycine (Sigma-Aldrich), which was incubated at RT for 5 min, then at 4°C for 15 min. Fixed cells were then collected by centrifugation at 3220 *g* for 10 min at 4°C, flash frozen on dry ice, and stored at -80 °C, to permit consistent processing of all samples in parallel.

Frozen pellets were thawed on ice, washed with 5 ml water, and centrifuged at 3220 *g* for 10 min. Pellets were then resuspended in 50 µl 30 mg ml^-1^ lysozyme, followed by incubation at 37°C for 30 min. Both 10% v/v sodium dodecyl sulphate (SDS; Fisher Bioreagents) and 10% v/v Triton-X (Sigma-Aldrich) were added, followed by 10 min RT shaking incubation, then 10 min incubation at 55°C. A 400 µl sample of the lysed cells was then combined with 130 µl of digestion mix, comprising NEB CutSmart buffer (NEB), 10% v/v Triton-X, and 1 U µl^-1^ of either *Nla*III (NEB; Illumina Hi-C library preparation), or *MluC*I (NEB; Pore-C library preparation). The chromatin was digested at 37°C for 3 h. The *Nla*III enzyme was then heat inactivated through a 65°C incubation for 20 min. The less stable *Mlu*CI enzyme did not require inactivation.

Digested chromatin was then ligated through the addition of 460 µL ligation mix, comprising ligation buffer (NEB), 0.1 µg µl^-1^ recombinant albumin (Promega), and 20 U µl^-1^ T4 DNA ligase (NEB). This mixture was incubated at RT for 1 h, followed by a 4°C incubation for 48 h. After ligation, the DNA-protein crosslinks were reversed through incubation of the mixture with 500 µl 20% Tween-20 (Sigma-Aldrich), 100 µl 10% v/v SDS, and 100 µl 20 µg µl^-1^ proteinase K (Sigma-Aldrich) at 65°C overnight. For the Pore-C library preparation, another round of protein denaturation was performed by adding 118 µl 20 mg ml^-1^ pronase (Sigma-Aldrich) and 235 µl 10% v/v SDS, followed by a 63°C incubation for 1 h.

Following de-cross-linking, DNA was purified with 2.5 ml of a chilled 25:24:1 ratio mixture of phenol:chloroform:isoamyl alcohol (Sigma-Aldrich), followed by separation by centrifugation at 3220 *g* for 30 min at 4°C. DNA was then precipitated from the aqueous phase of each sample using 20 µl 5 M NaCl (Sigma Aldrich), 200 µl 3 M sodium acetate (EMD Millipore Corporation) and 6 ml isopropanol (Sigma-Aldrich). These mixtures were incubated at -80°C overnight, then centrifuged at 3220 *g* for 30 min at 4°C. The resulting pellets were washed with 1 ml of chilled 70% ethanol (VWR Chemicals BDH), followed by a second centrifugation (3220 *g*, 15 min, 4°C). All remaining ethanol was removed through pipetting and drying at 65°C in a heat block for 15 min. The DNA was the resuspended in 100 µl nuclease-free water.

To enrich for longer molecules likely to contain multiple ligated DNA fragments, a 100 µl sample of each preparation was mixed with 50 µl of KAPA Pure Beads (Roche) through vortexing, followed by incubation together at RT for 10 min. The beads were then magnetically captured, and the supernatant discarded. The beads were washed twice using 200 µl 80% ethanol and dried at RT for 4 min. DNA was then eluted from the beads through a 10 min incubation in 100 µl of pre-heated nuclease-free water at 37°C. The concentration and size distribution of the DNA molecules was checked using an Agilent 2200 Tape Station (Agilent Technologies) and NanoDrop 1000 (Thermo Fisher).

### DNA sequencing

Illumina Hi-C data were generated at the Wellcome Sanger Institute. Libraries were prepared using standard approaches [70], enriching for a fragment size of approximately 450 bp. Libraries corresponding to the three biological replicates for each of the four variants were then combined into a 12-plex library, which was sequenced on a single lane of the Illumina Novaseq 6000 platform. This generated 152 nt paired end reads.

Pore-C data were generated at Imperial College London using the native 12-plex barcoding library preparation kit (Oxford Nanopore Technologies, ONT; product SQK-NBD114.24) to process ∼400 ng of DNA from each of the three biological replicates for each of the four variants. Each DNA sample was individually treated to ensure fragments were blunt ended with the NEBNext FFPE DNA Repair Mix (NEB) and the NEBNext Ultra II End Repair/dA-tailing Module (NEB). Samples then underwent native barcode ligation with NEB Blunt/TA Ligase Master Mix, before being pooled together. The NEBNext Quick Ligation Module (NEB) was then used to attach sequencing adapters. The completed libraries were then purified from the mixture using the short fragment buffer to obtain DNA fragments of all sizes. The concentration of the final DNA library was measured on a Qubit 4 fluorometer (Thermo Fisher). Two flow cells (ONT product FLO-MIN114) were each used to sequence 15 fmol of the final library using the MinION platform, recording data with a minimum quality score of 9 using high-accuracy basecalling with Guppy version 6.4.6 (ONT).

### Bioinformatic processing

The Illumina Hi-C data were processed using version 2.1.0 of the hic workflow [71] implemented using Nextflow version 24.02.0 [72]. This workflow filtered the Illumina reads using FastQC [73], then mapped reads to the appropriate reference genome (Table S1) using bowtie2 [74], and generated raw contact maps using iced, as implemented within the HiC-Pro pipeline [75]. Genome-wide contact maps between loci were then generated at resolutions of 100, 500, 1000, 5000, 10000 and 50000 bp using cooler [76]. A quality control report was then generated with MultiQC [77], with additional visualisation files produced with juicer [78].

The Pore-C data were processed using version 1.1.0 of the wf-pore-c workflow [79] implemented using Nextflow version 23.10.1 [72]. This workflow mapped reads to the appropriate reference genome (Table S1) using minimap2 [80] with the non-default settings “-x map-ont -k 7 -w 5”. These alignments were then processed with Pore-c-py [79] and pairtools [39] to generate raw contact maps, which were processed with cooler [76] and juicer [78], as for the Hi-C data.

### Differential analysis of contact matrices

Analyses of differential contacts used matrices calculated at a resolution of 10 kb, with all self interactions along the matrix diagonals removed, and the *ivr* and *tvr* loci excluded, as they contained known rearrangements.

The diffHiC package [43] was applied using the recommended workflow [81], with modifications to parameter settings as appropriate for these datasets. Loci with fewer than five contacts across all replicates in each comparison were first removed, as there was low power to identify differential contacts at such positions. The counts across the filtered dataset were then normalised across replicates within each comparison between variants using a cyclic LOESS process, run for ten iterations. The biological coefficient of variation was then calculated across different contact densities, which informed the dispersion of the negative binomial generalised linear model that was fitted to the data. This used a tagwise, rather than trended, dispersion calculation, as there was not a strong relationship between variance and contact densities in some datasets. Differential contact densities between loci in the two variants were then identified from this model using quasi F-tests. The resulting *p* values were subject to a Benjamini-Hochberg correction for multiple testing to generate *q* values.

Similarly, analysis with the multiHiCcompare package [44] used a modified version of the recommended workflow [82]. This started by first both removing any loci for which the minimum frequency across replicates was zero. The remaining data were normalised using a cyclic LOESS process run for ten iterations. Significant differences were inferred through using a negative binomial exact test, as implemented in edgeR [83], with a Benjamini-Hochberg correction for multiple testing.

### Statistical modelling of contact densities

The 10 kb resolution contact matrices were analysed using a generalised linear mixed effects model with the structure:

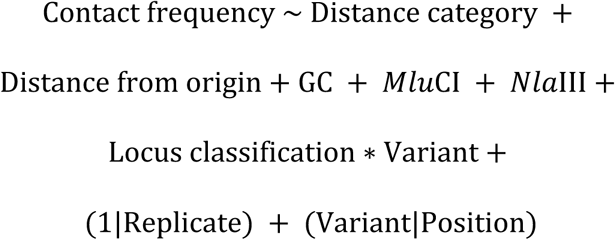

The terms all represent fixed effects, except those with the ‘|’ notation, which denotes a grouping variable for a random effect. Asterisks indicate an interaction between fixed effects. A zero-inflated logarithmic Poisson link function was used to fit the model with the glmmTMB package [84]. Parameter estimates reproducibly converged using the default settings. However, attempts analyse the data using an equivalent model with a negative binomial link function, which would account for any overdispersion in the data, failed to converge on robust parameter estimates. Analysis of parameter estimates used the sjPlot package [85], model fits were assessed with the performance package [86], and data processing used the tidyverse packages [87].

Each interaction was categorised by the distance between the loci: <12.5 kb (ie., neighbouring 10 kb loci); 12.5-25 kb; 25-50 kb, and >50 kb. Only loci within 100 kb of each other were considered, to ensure parameter estimation was feasible, and avoid the model fitting primarily to the noisy, sparse interactions between distant loci.

Loci were also classified based on the annotation of the corresponding reference genomes. In both genomes, the four rRNA operons were annotated, as the difficulty of mapping to these relatively GC-rich repeats reduces the detection of contacts involving these loci. In RMV7, four MGEs were annotated: PRCI*_dnaN_*, PRCI*_uvrA_*, ICE*_rpsI_*, and Tn*916* [12]. In RMV8, three MGEs were annotated: the prophage remnant CIPhR, prophage ɸRMV8 and PRCI*_malA_* [18]. These classifications were fitted separately for each variant, to enable intervariant differences to be inferred at these loci.

As the DNA-protein crosslinks were fixed in mid-exponential phase, the detected contact frequencies were higher near the origin of replication, as some cells would have initiated, but not completed, chromosomal replication. Hence the number of detected contacts was expected to change with distance from the origin.

The additional fixed effects analysed the properties of each DNA locus, which were calculated using biopython [88]. The effect of the distribution of *Nla*III and *Mlu*CI sites was accounted for through calculating the mean number of restriction sites across the two interacting loci, and dividing this value by ten. The GC content was calculated as the mean proportion of bases in the two interacting loci, which was scaled upwards by a factor of ten. Similarly, the distance to the origin was calculated as the proportion of the genome between the site and the origin of replication, and this value was scaled upwards by a factor of ten when included in the model. These adjustments were necessary for numerical stability of the model fitting process.

The identity of the biological replicate was included as a random effect, to correct for differences in the amount of sequence data generated for each dataset. A second random effect estimated the corrected level of contact frequency for each locus in the dominant variant, and the difference in binding to each of these loci in the rare variant.

## Data Availability

All raw genetic data used in this study are available from the ENA with the accession codes listed in Table S1 and S2. Processed files generated by bioinformatic pipelines are available from FigShare (https://figshare.com/projects/Hi-C_and_Pore-C_data_for_pneumococcal_epigenetic_variants/227616). Experimental results and sample information are available from Github (https://github.com/nickjcroucher/RMV_PoreC).

## Code Availability

All code used in this analysis is available from Github (https://github.com/nickjcroucher/RMV_PoreC).

## Acknowledgements

We thank Prof. Juanma Vaquerizas for helpful discussions about Hi-C data generation and analysis, and the Bespoke team at the Wellcome Sanger Institute for generating the Hi-C sequencing libraries.

## Supporting Information Legends

**Supporting Information File 1** This file contains Figures S1-S25 and Tables S1-S4, and their associated legends.

