## Supplementary Figures and Tables for "Pore-C sequencing identifies episome-driven chromosome conformation perturbations differentiating pneumococcal epigenetic variants"

**Supplementary Materials:**

Figures S1-S25

Tables S1-S4

**Supplementary Figures**

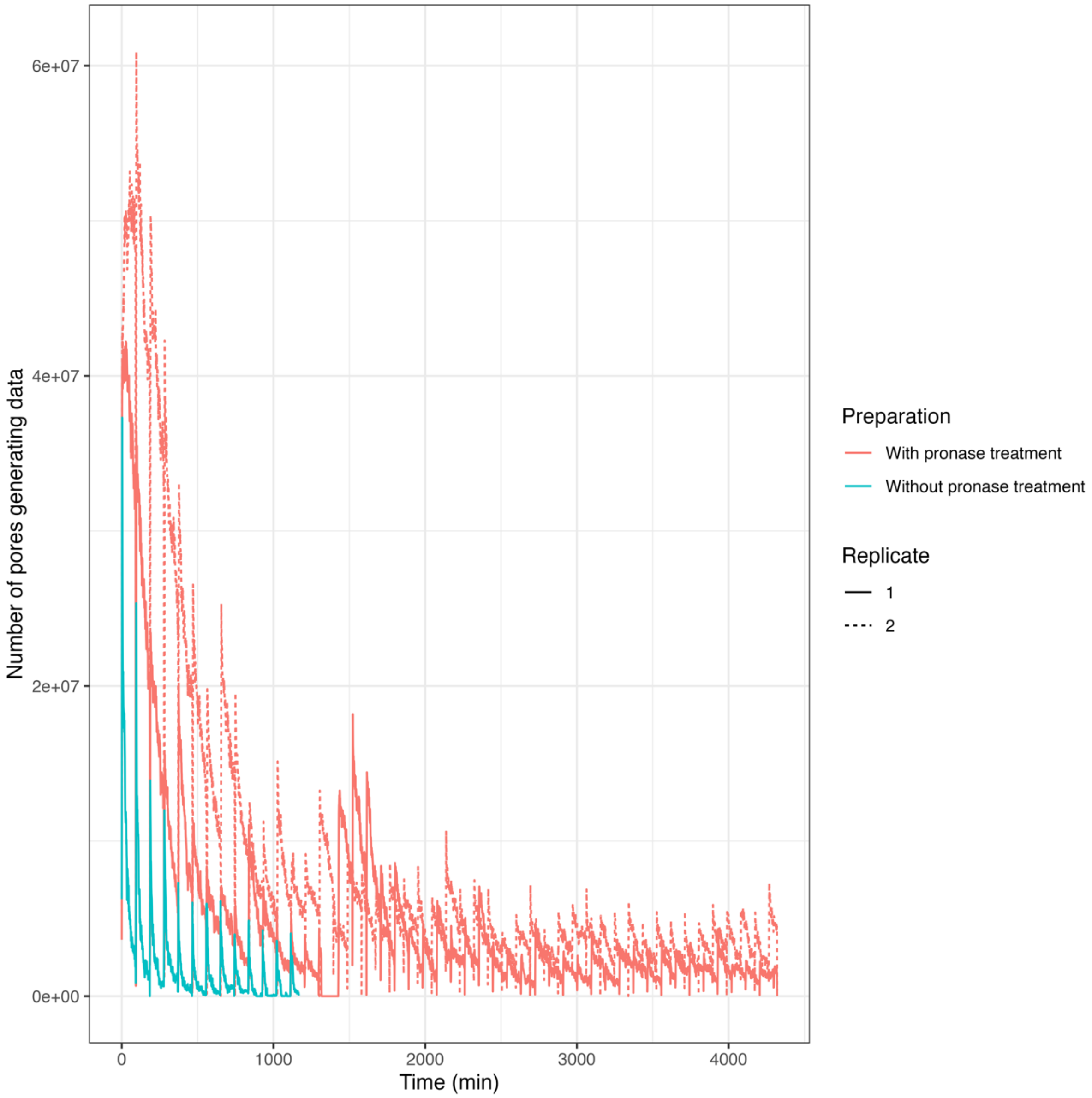


**Figure S1** A comparison of the data generated from Pore-C libraries produced using different protocols. The line plots show the number of nanopores on a flow cell generating sequence data over the course of three sequencing runs. This declined over time as pores became blocked by DNA that remained cross-linked to proteins. The MinION device attempts to clear such blockages through regular transient voltage reversals, resulting in the jagged appearance of the curve. The lines are coloured according to whether the sequencing library was prepared using de-cross-linking with proteinase K only (one replicate), or proteinase K and pronase (two replicates). This demonstrated the additional pronase treatment substantially reduced the rate of pore blocking, resulting in an increased yield of sequence data.

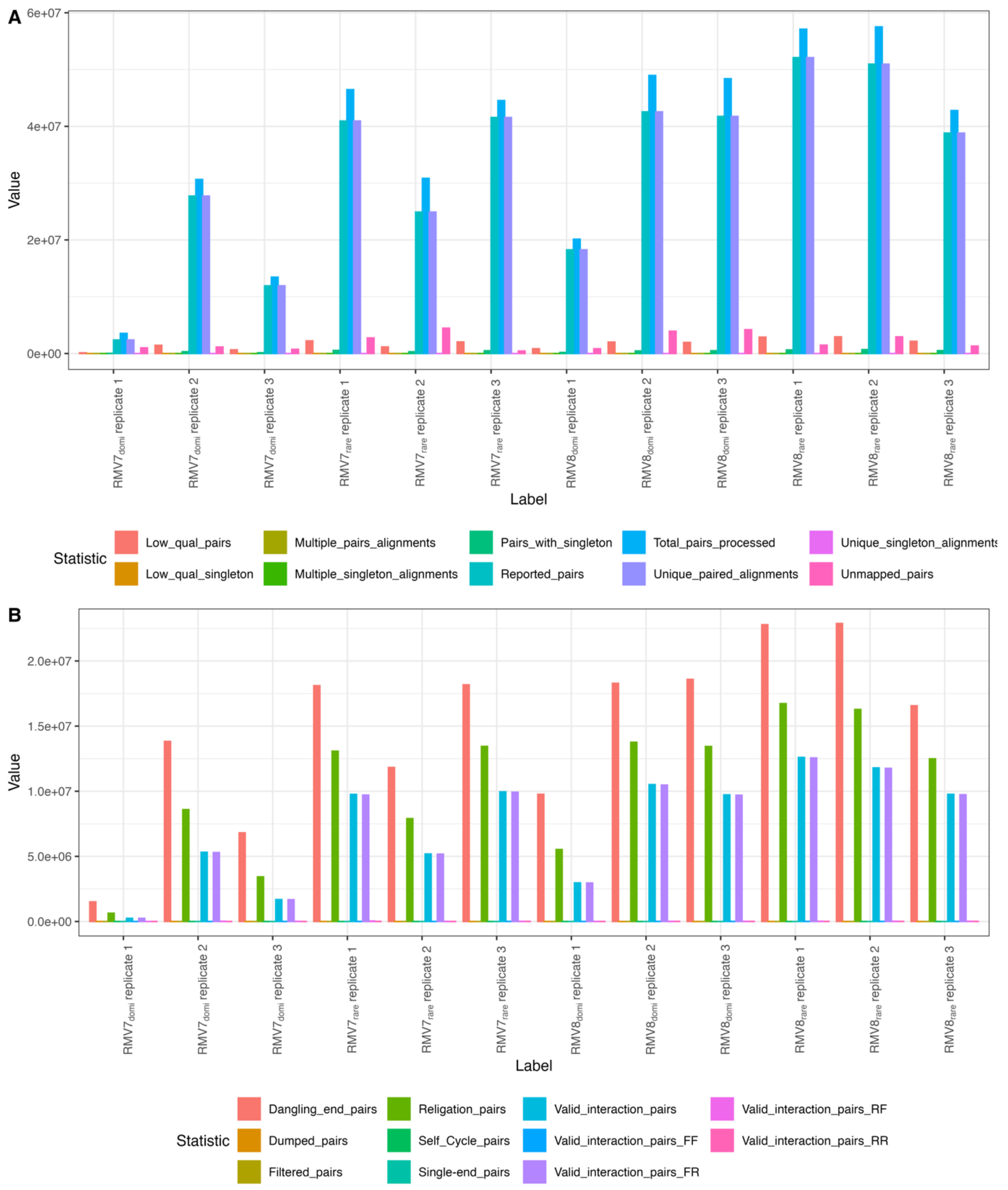


**Figure S2** Statistical description of the Illumina Hi-C data by the HiC-Pro pipeline. **(A)** Analysis of paired read alignment to the reference genome. This shows the majority of processed pairs were successfully mapped to the genome as unique, paired alignments. **(B)** Analysis of the aligned pairs. Most reads were split between three categories. The “dangling end pairs” correspond to reads from the ends of the same restriction fragment, which represents a failure to ligate a DNA molecule into a longer concatemer. The “relegation pairs” correspond to either to DNA molecules that were not digested, or neighbouring fragments that were ligated back together into their original form. The “valid interaction pairs” correspond to pairs that allow for a digestion and relegation to be inferred. Almost all of these pairs have the expected forward-reverse relative orientation of the reads.

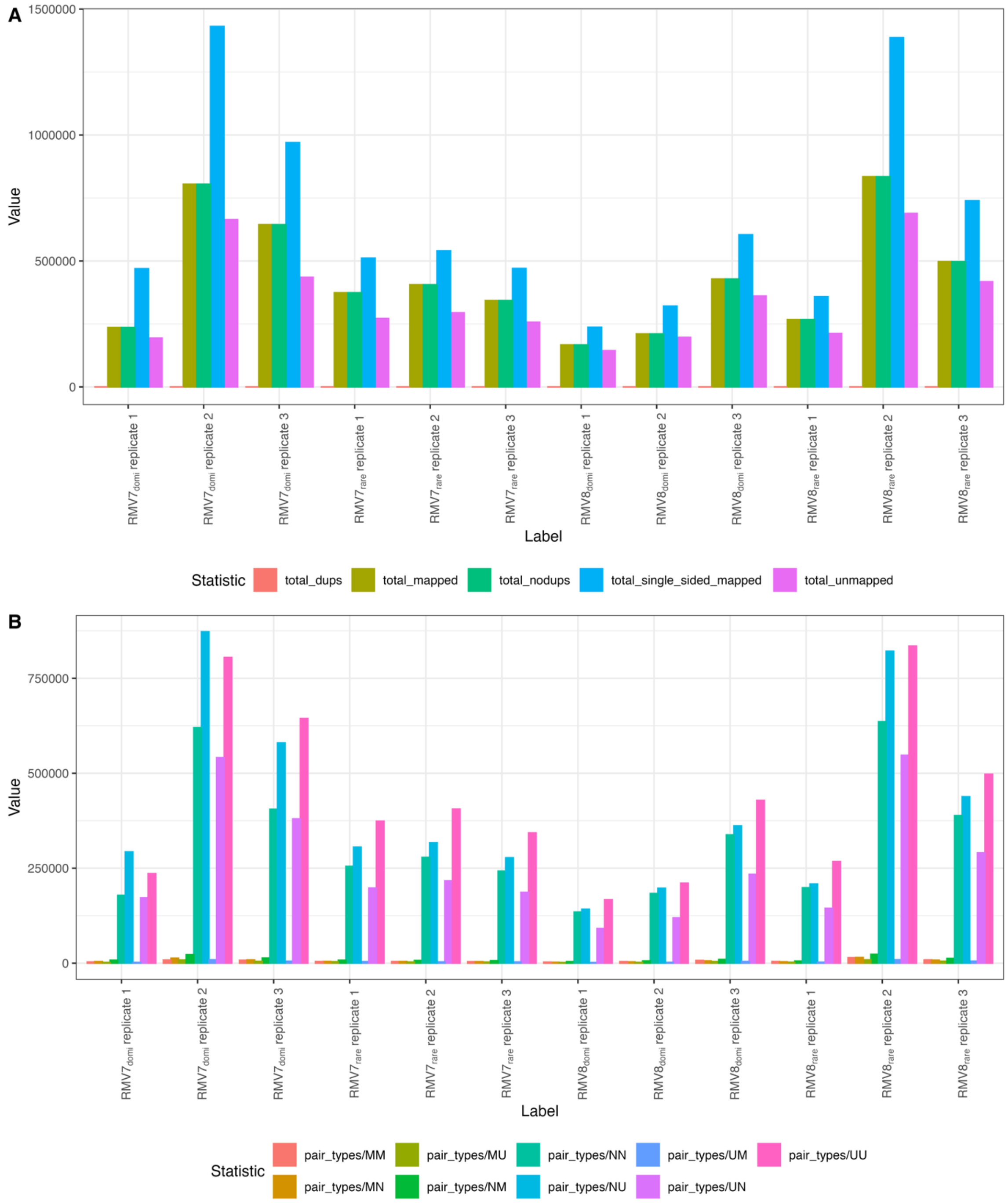


**Figure S3** Analysis of the Pore-C mapping data generated by pairtools. **(A)** Processing of sequence reads, split into restriction fragments. The modal categories comprised pairs in which only one of the two sequences could be mapped to the reference genome. Nevertheless, there were more pairs of sequences that could both be mapped to the reference than pairs of sequences in which neither fragment mapped to the reference. **(B)** Details of read mapping. All pairs were classified by the mapping status of the constituent reads. The unmapped reads from (A) were divided between categories in which at least one read was not able to be mapped to the reference genome (N), or mapped to multiple sites (M). The single-sided mapped reads were split into the four categories in which only one read could be uniquely mapped (U).Yet many read pairs consisted of paired fragments that were both uniquely mapped (UU), enabling inference of contacts across the genome.

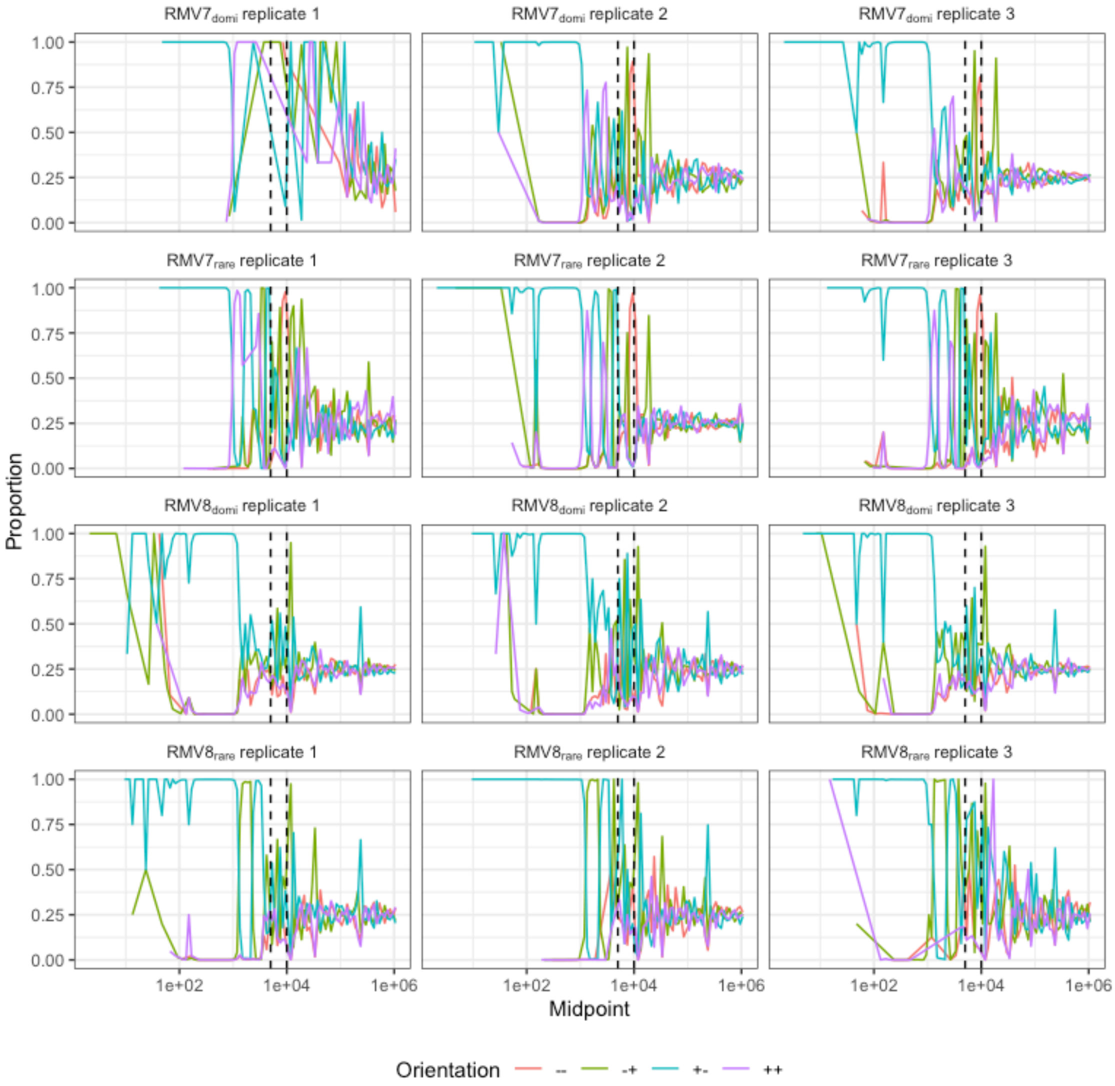


**Figure S4** Line plots showing the mapping orientations of read pairs from the Illumina Hi-C datasets at different separations. The read pairs were categorised into 100 bins, based on the distribution of logarithmically-scaled distances between the mapping locations of the pair members. For each bin, the proportions of read pairs mapping in the four possible orientations (both to the positive strand; both to the negative strand; the forward read mapping to the positive strand and the reverse read mapping to the negative strand; or the forward read mapping to the negative strand and the reverse read mapping to the positive strand) were calculated. The different orientations are represented by the colour of the lines. As Illumina read pairs are generated by sequencing initiated from each end of a DNA molecule, reads generated from undigested templates are expected to map to different strands of the genome (i.e. have a +/- or -/+ orientation). These orientations dominate over short distances, suggesting read pairs mapping a short distance from one another were generated from uncut genomic DNA. The dashed lines represent estimates of the convergence distance, at which digestion and religation is sufficiently frequent for the four orientations to occur at an approximately equal frequency of ~0.25. Each panel shows the data from an individual replicate.

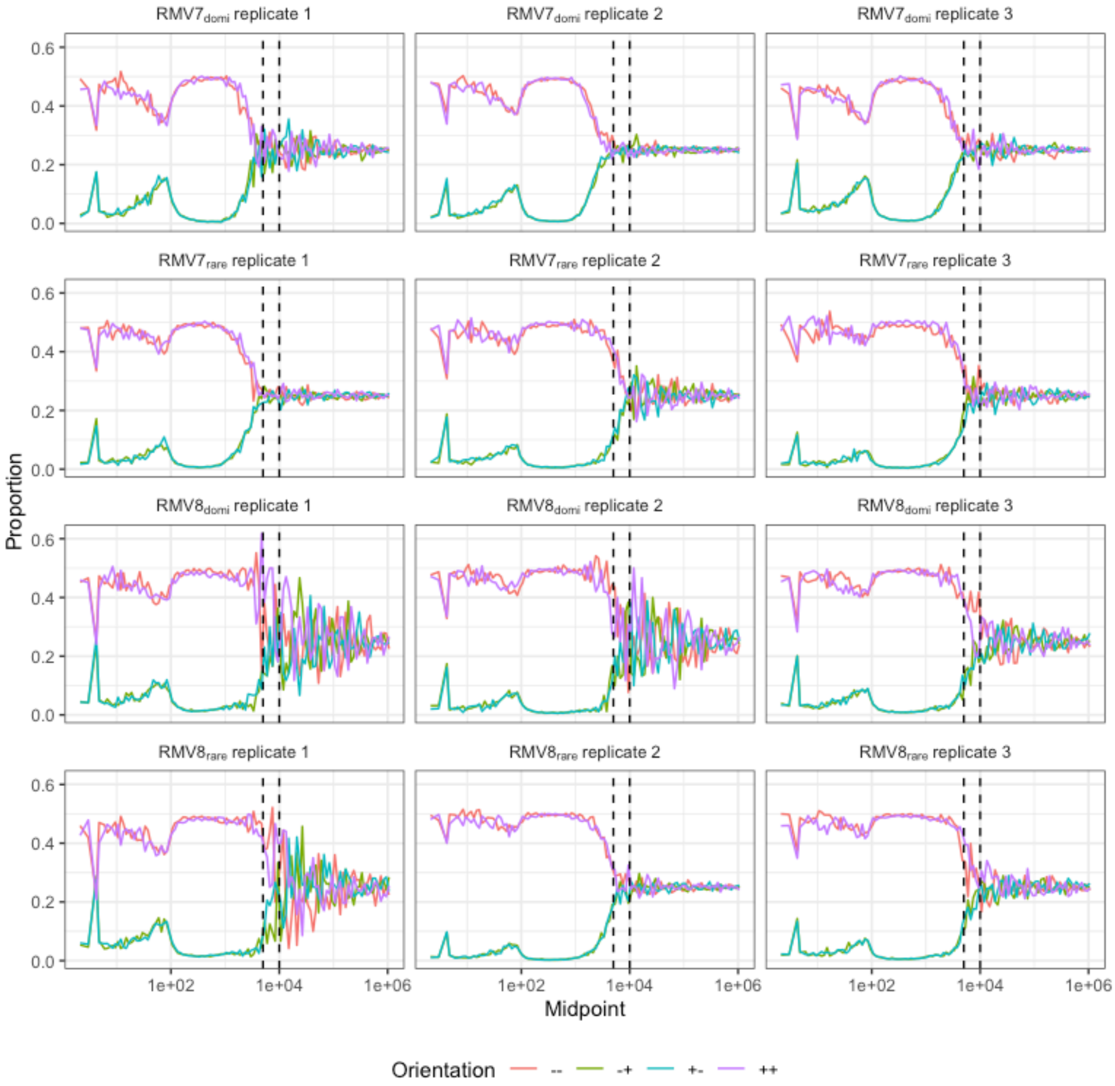


**Figure S5** Identification of the convergence distance in the Pore-C data. Data are shown as in Fig. S4. As Nanopore reads are generated as continuous sequence following initiation at one end of a molecule, data generated from an undigested template is expected to have a +/+ or -/- orientation. Therefore these orientations dominate over short distances, in contrast to the analyses shown in Fig. S4.

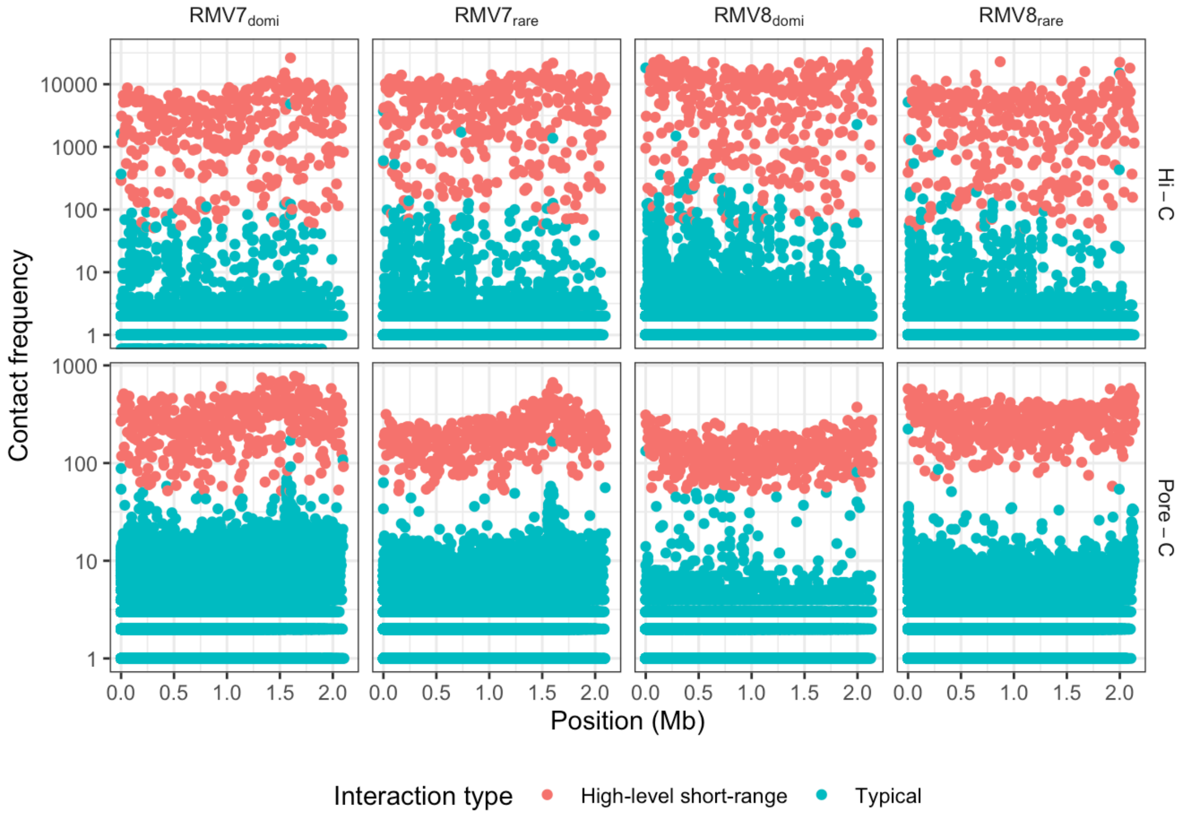


**Figure S6** Spatial distribution of the high-level short-range interactions between loci highlighted in Fig. 1B. The absence of any spatial clustering of the short-range highly-interacting loci suggested they could not be explained by a process localised to a specific region, and therefore were most parsimoniously explained as artefactual products resulting from incomplete digestion of DNA.

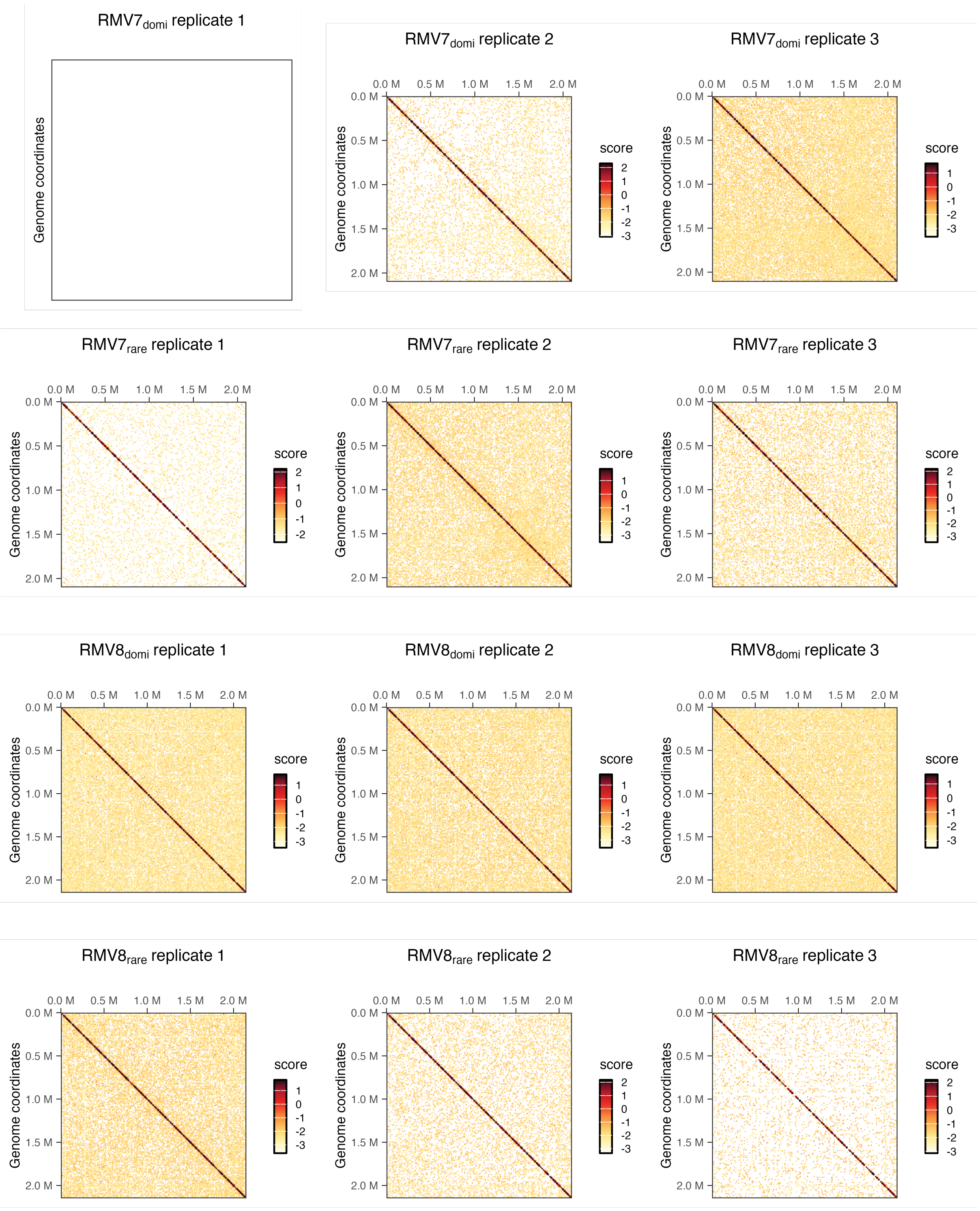


**Figure S7** Contact matrices for individual Illumina Hi-C replicates, calculated at a resolution of 10 kb. Insufficient long-range interactions were inferred from the first sample extracted from *S. pneumoniae* $\text{RMV7}_{\text{domi}}$ to generate a matrix. Each other matrix is symmetrical, with both the horizontal and vertical axes representing the length of the genome. Each cell is coloured to represent the frequency of interactions between the corresponding loci. Keys are provided for each replicate individually, to adjust the visualisation to the amount of data generated.

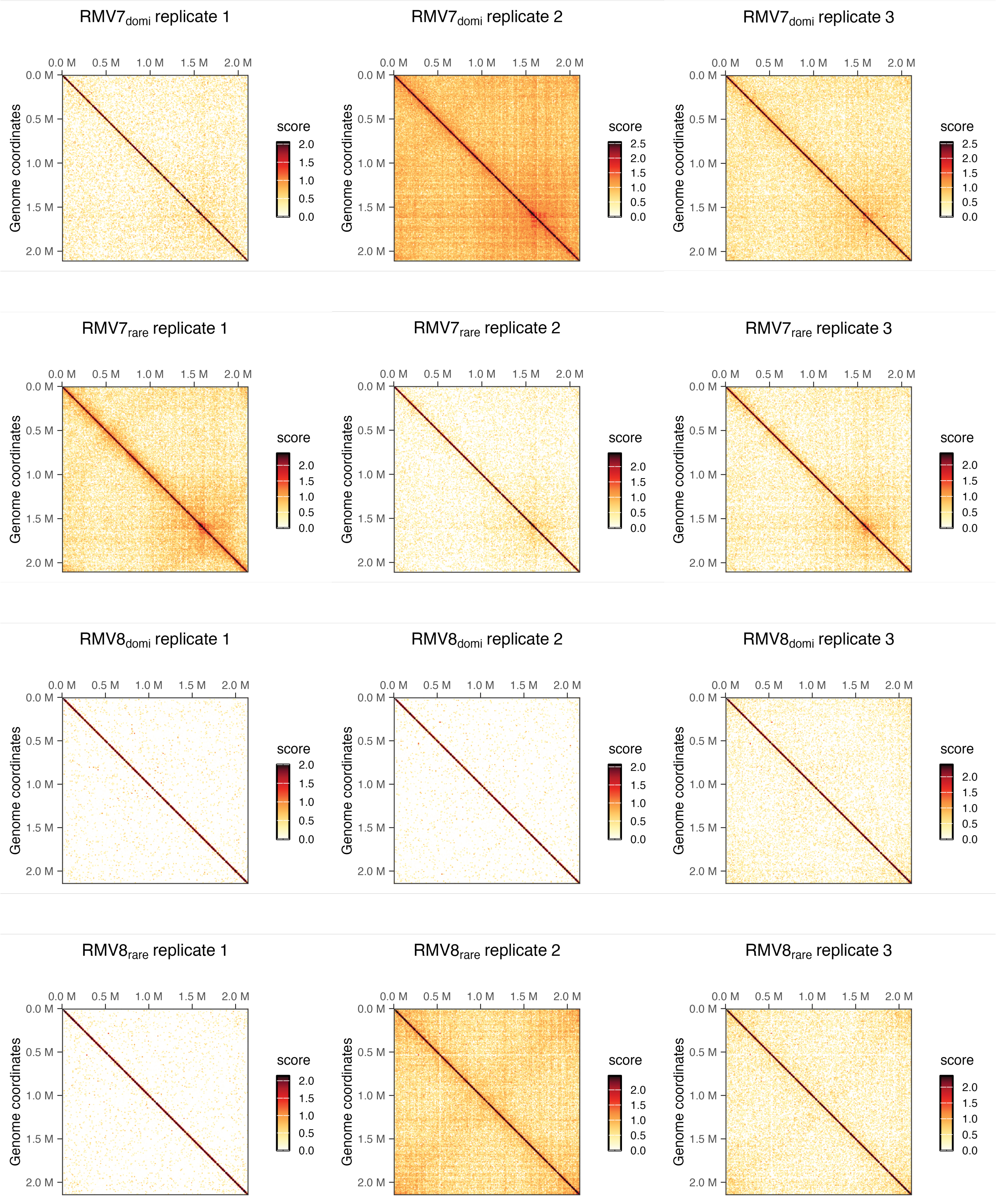


**Figure S8** Contact matrices for individual Pore-C replicates, calculated at a resolution of 10 kb. Data are shown as in Fig. S4.

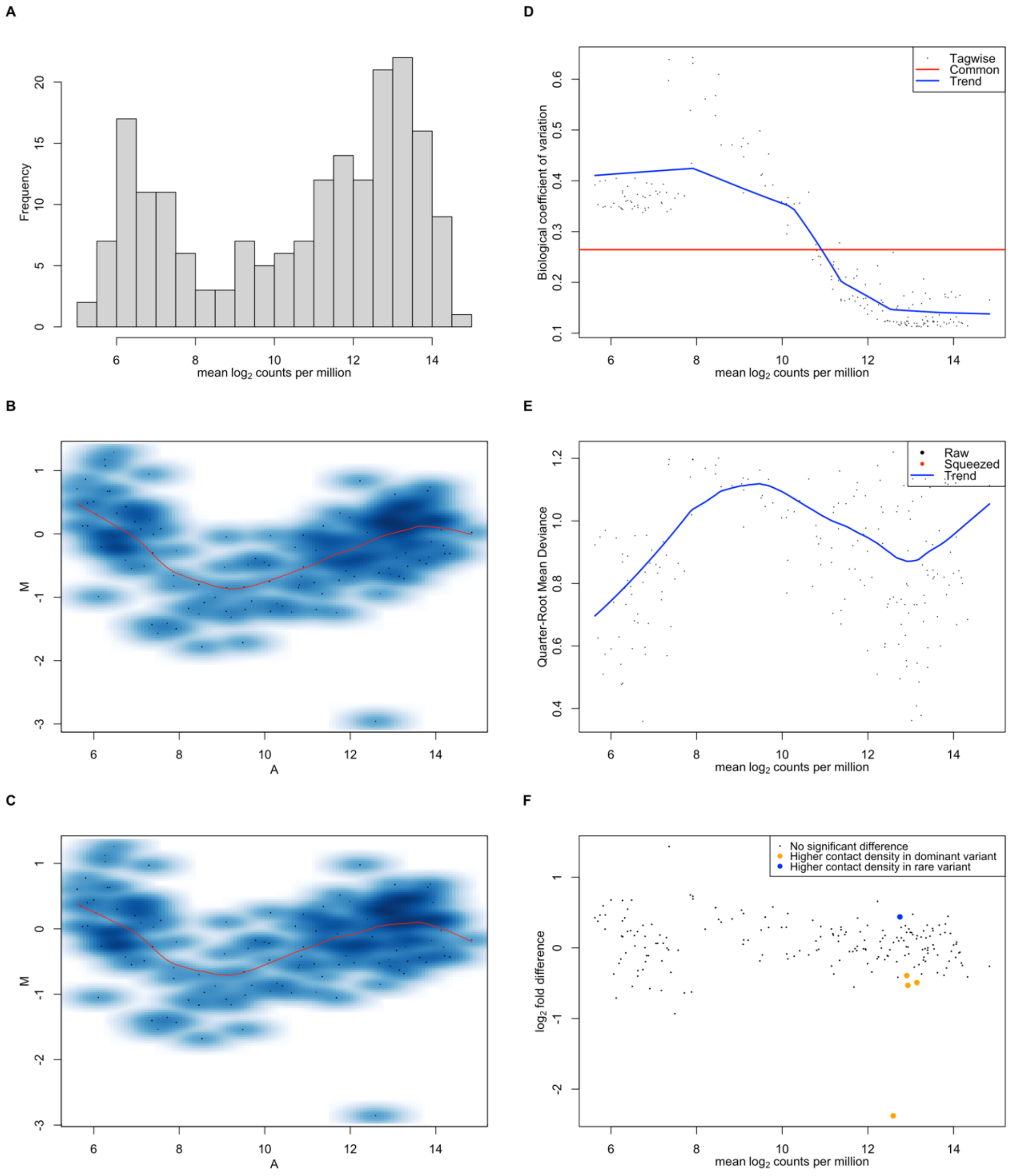


**Figure S9** Processing of Hi-C data from *S. pneumoniae* RMV7 for analysis with diffHiC. All analyses were conducted using a locus size of 10 kb. **(A)** Histogram showing the distribution of contact frequencies across all loci and replicates as mean $\log_2$ counts per million reads. **(B)** MA plot comparing the mean $\log_2$ counts per million reads (denoted A) against the ratio of $\log_2$ counts per million reads in the rare variant relative to the dominant variant (denoted M) for a representative pair of biological replicates. The density of points is represented by the blue shading. The LOESS regression line is shown in red. **(C)** MA plot of the same data after normalisation. **(D)** Relationship between the biological coefficient of variation, relating to the dispersion parameter of the negative binomial distribution used to fit the generalised linear model, and the mean $\log_2$ counts per million reads. **(E)** Relationship between the quasi-likelihood dispersion and the mean $\log_2$ counts per million reads estimated by fitting the generalised linear model to the contact frequency data. **(F)** MA plot of normalised data, highlighting loci found to have a significantly higher density of contacts in the two variants.

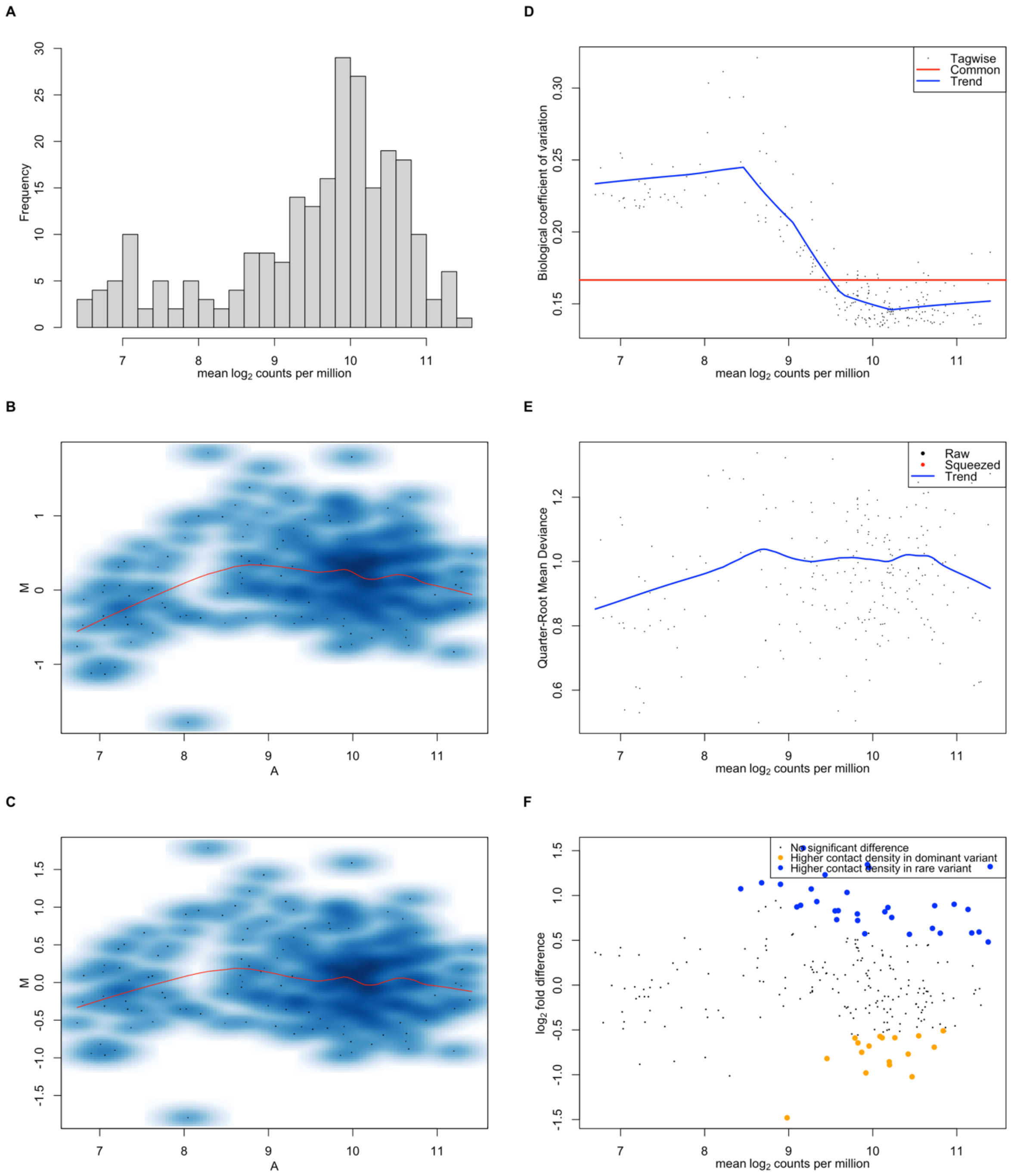


**Figure S10** Processing of Pore-C data from *S. pneumoniae* RMV7 for analysis with diffHiC. All analyses were conducted using a locus size of 10 kb. Data are shown as in Fig. S9.

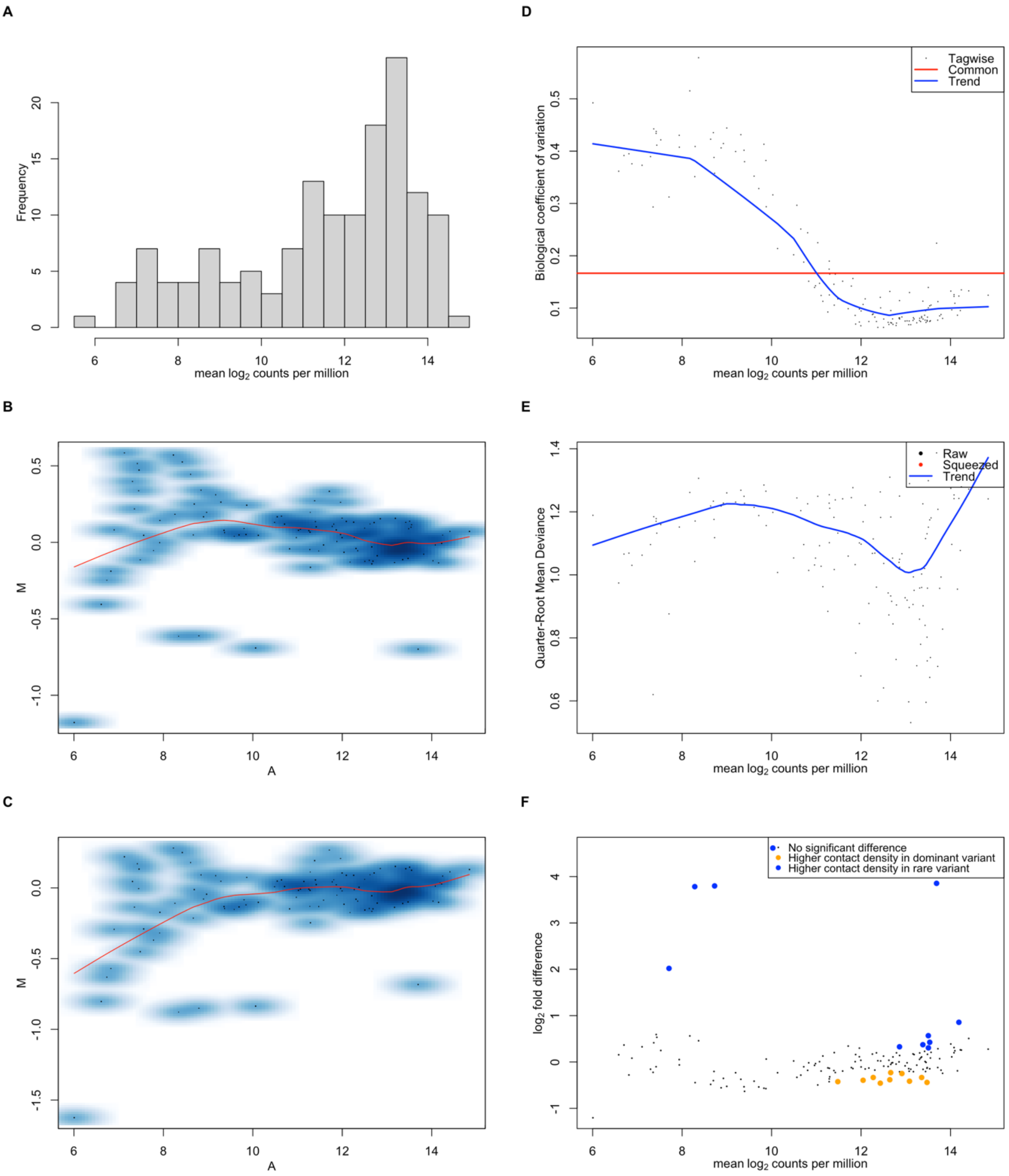


**Figure S11** Processing of Hi-C data from *S. pneumoniae* RMV8 for analysis with diffHiC. All analyses were conducted using a locus size of 10 kb. Data are shown as in Fig. S9.

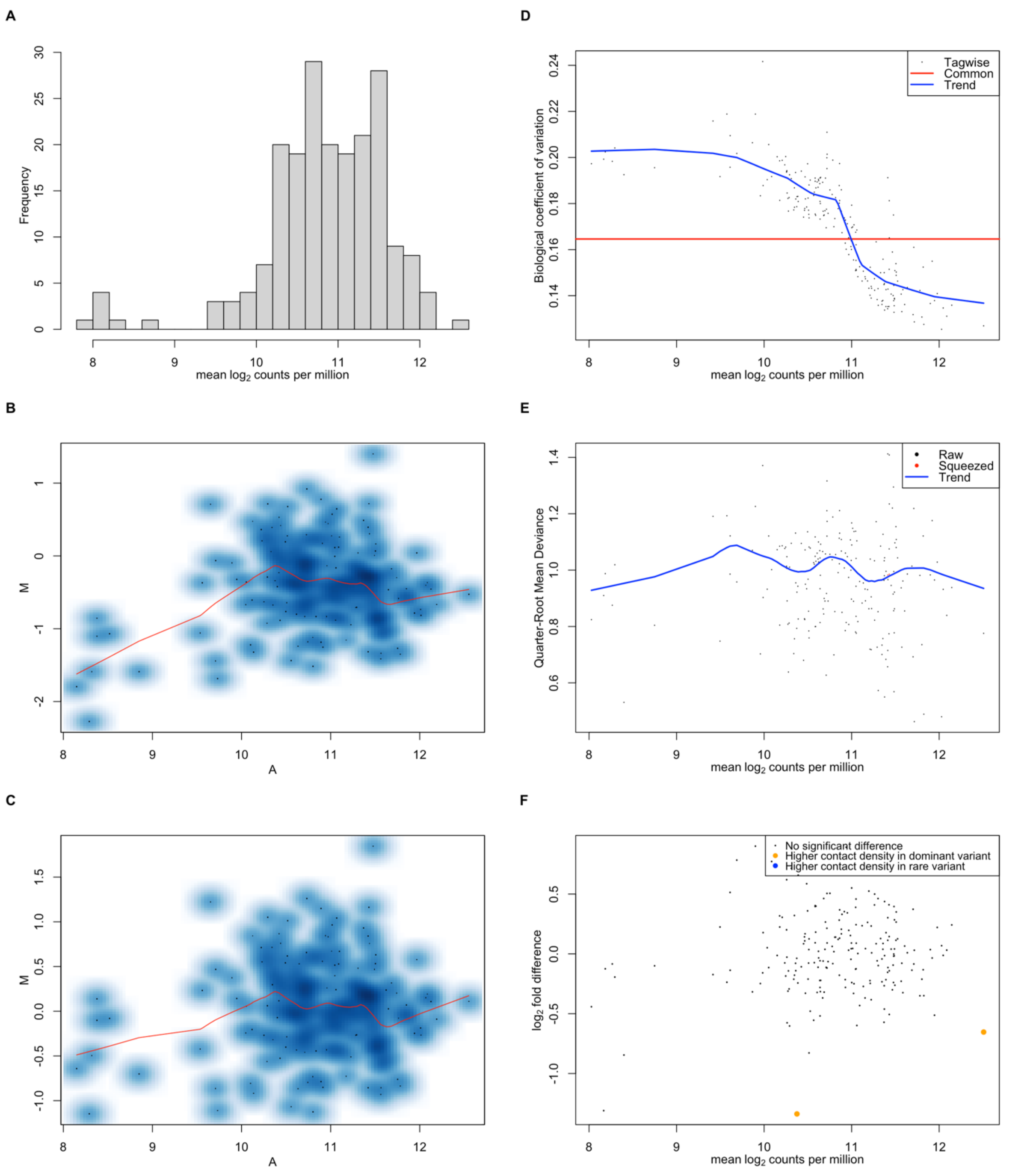


**Figure S12** Processing of Pore-C data from *S. pneumoniae* RMV8 for analysis with diffHiC. All analyses were conducted using a locus size of 10 kb. Data are shown as in Fig. S9.

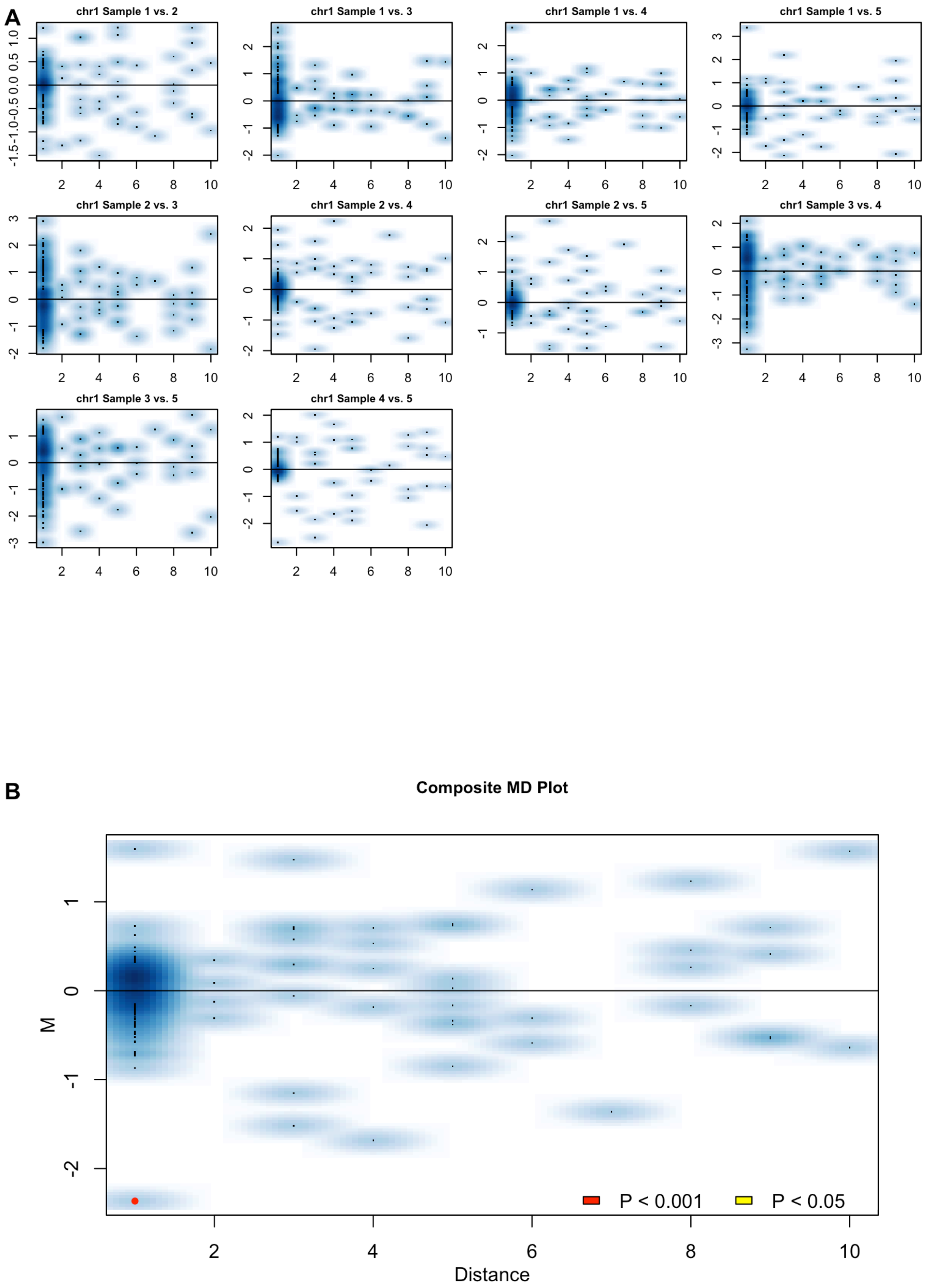


**Figure S13** Processing of Hi-C data from *S. pneumoniae* RMV7 for analysis with multiHiCcomparison. All analyses were conducted using a locus size of 10 kb. **(A)** MD plots for each pairwise comparison between replicates after joint normalisation by a cyclic LOESS process. Each plot shows the ratio of $log_2$ counts per million reads in the one replicate relative another (denoted M), relative to the distance between the interacting loci (denoted D), enumerated as the difference between the indices of the loci. **(B)** Composite MD plot summarising the output of the test for differential contact intensities across all replicates. The position of points on the vertical axis represents the $\log_2$ fold difference between the variants, with points higher on the axis when the contact density is higher in the rare variant relative to the dominant variant. Points are coloured according to the categorisation of their false discovery rate (i.e. *q* value), calculated after a Benjamini-Hochberg correction for multiple testing.

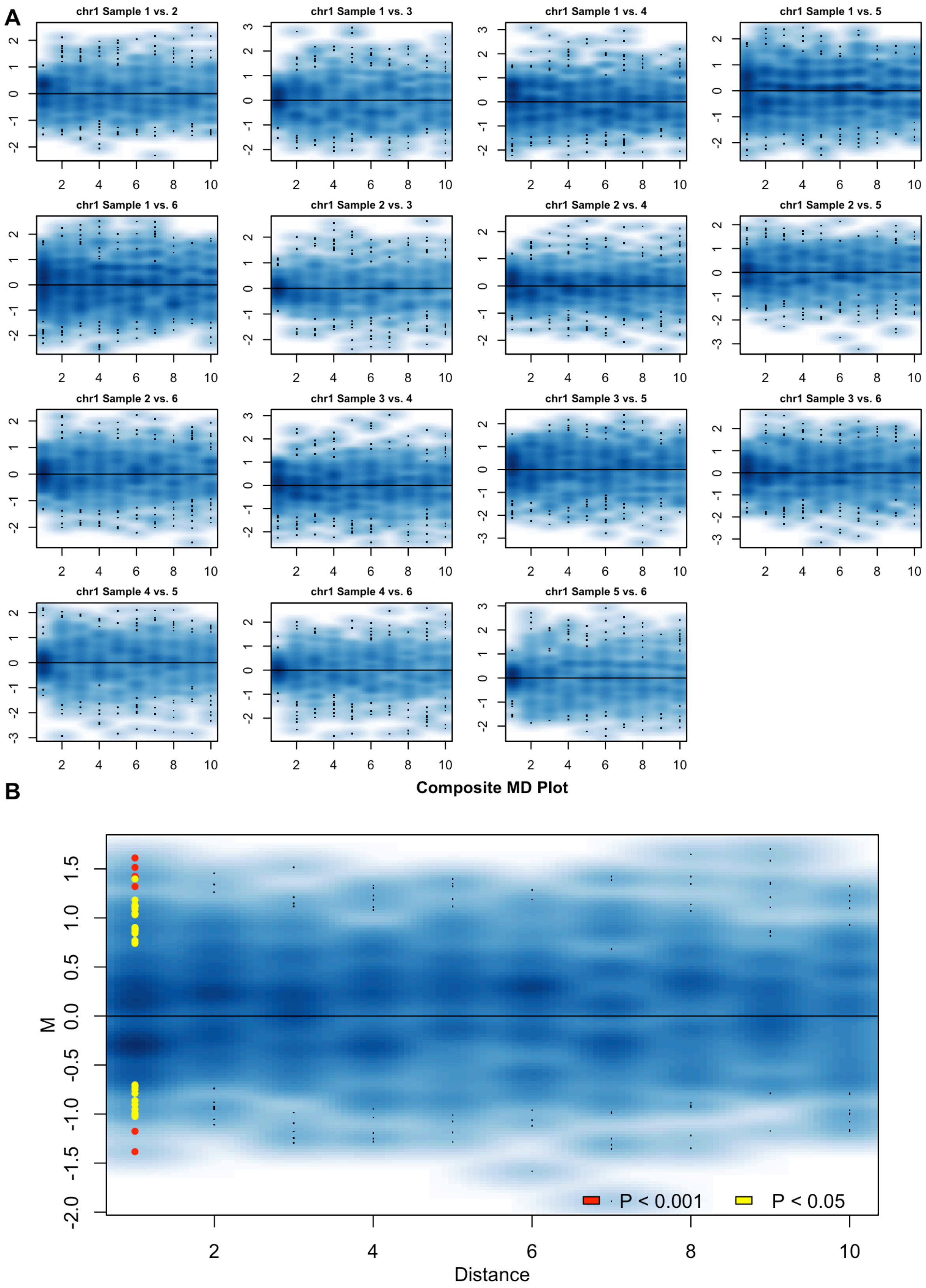


**Figure S14** Processing of Pore-C data from *S. pneumoniae* RMV7 for analysis with multiHiCcomparison. All analyses were conducted using a locus size of 10 kb. Data are shown as in Fig. S13.

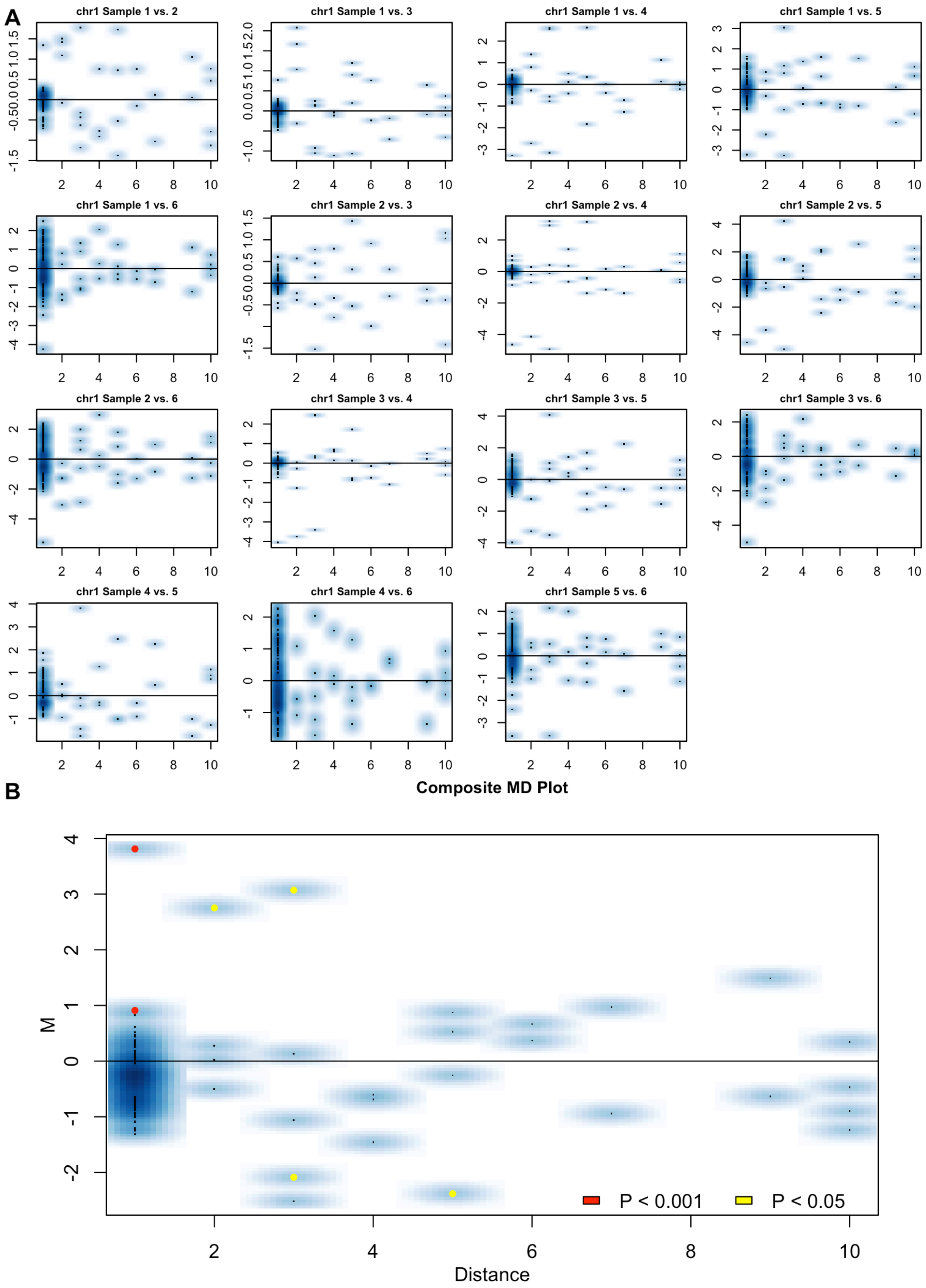


**Figure S15** Processing of Hi-C data from *S. pneumoniae* RMV8 for analysis with multiHiCcomparison. All analyses were conducted using a locus size of 10 kb. Data are shown as in Fig. S13.

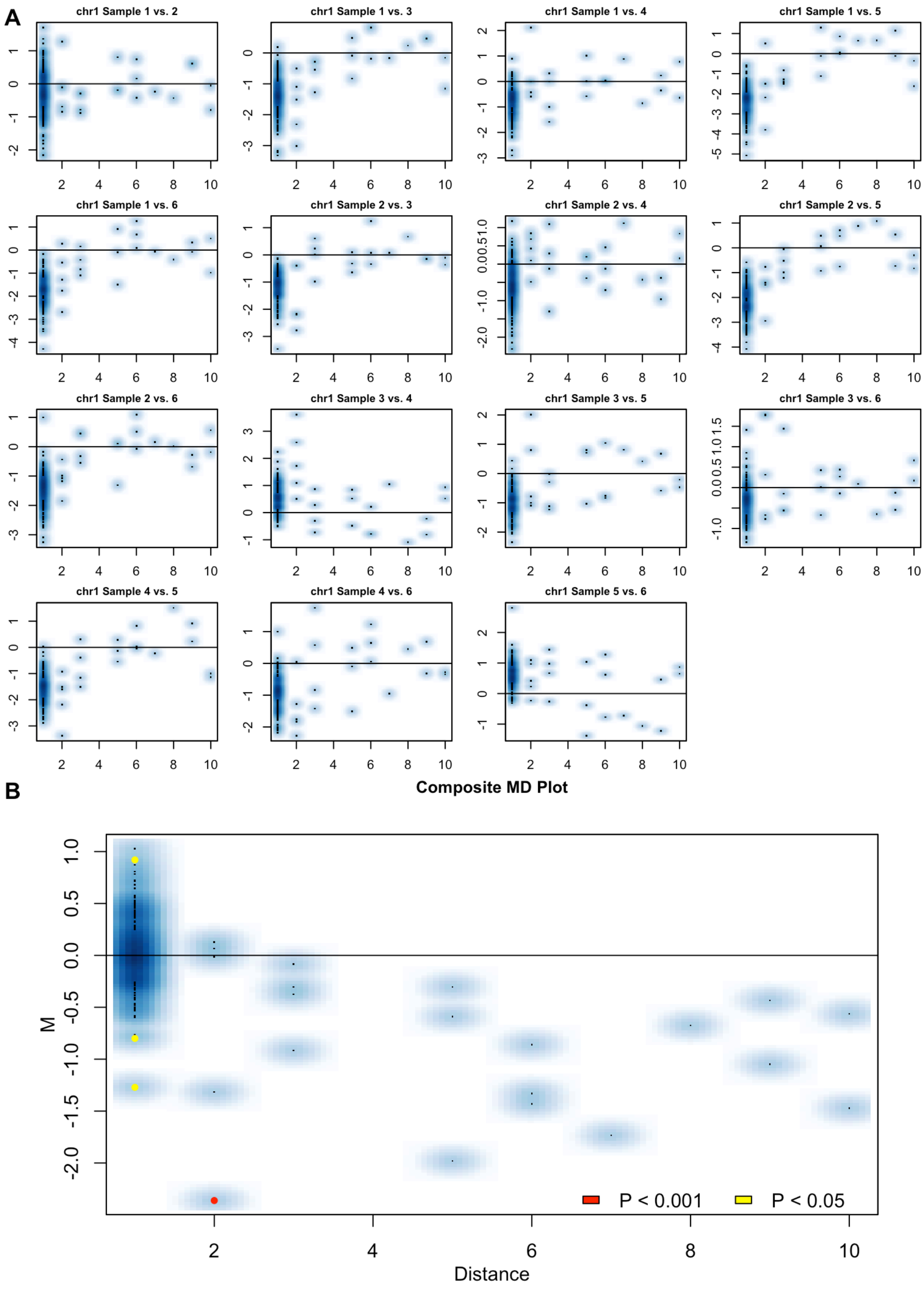


**Figure S16** Processing of Pore-C data from *S. pneumoniae* RMV8 for analysis with multiHiCcomparison. All analyses were conducted using a locus size of 10 kb. Data are shown as in Fig. S13.

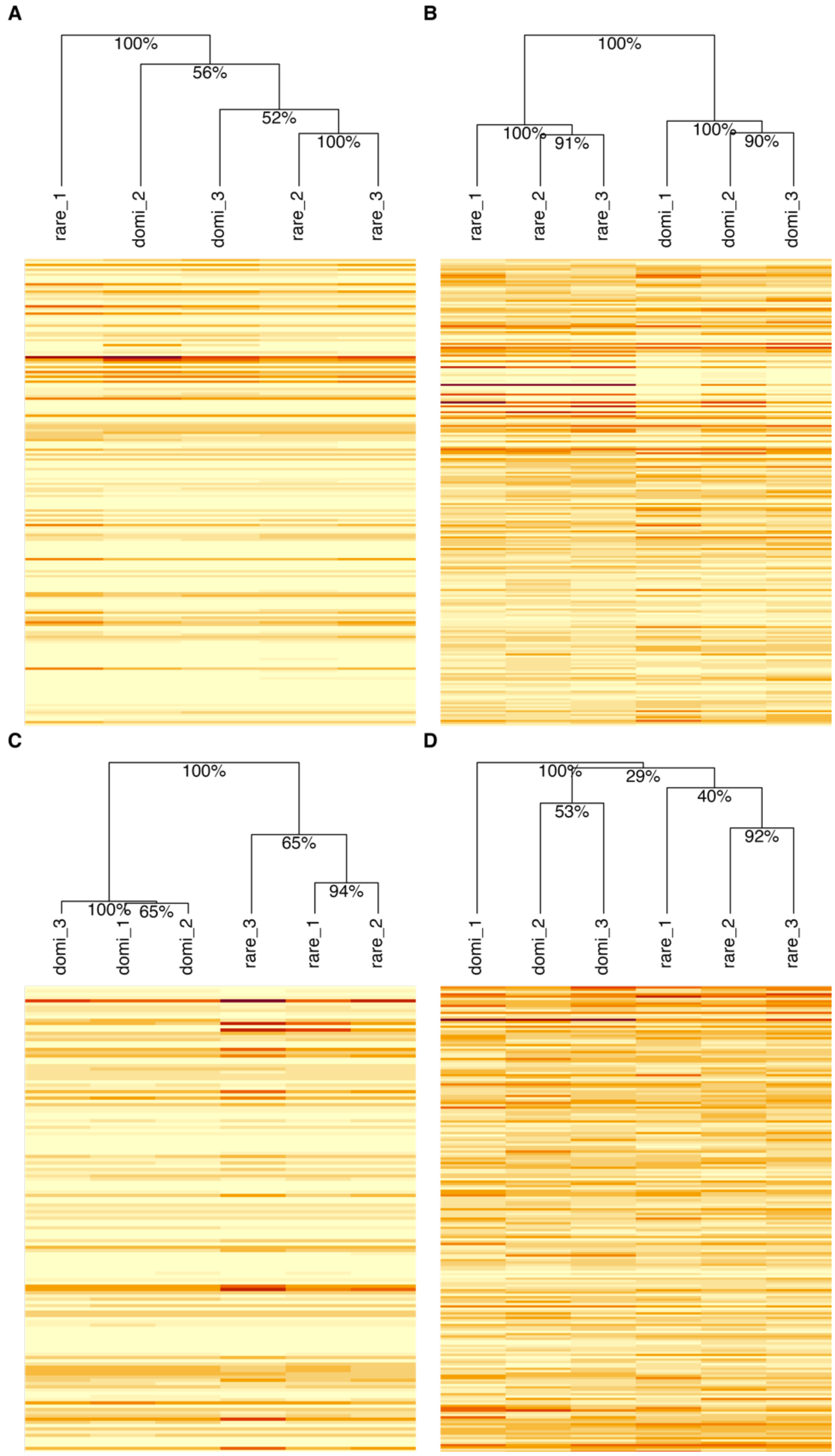


**Figure S17** Heatmaps comparing the contact frequency patterns between variants. Each column is a different replicate dataset from either the dominant (domi) or rare variant. Each row represents a locus. The colour of each cell represents the intensity of contacts at a locus in a replicate following normalisation as part of the diffHiC analyses. The variants are clustered based on their similarity, as shown by the dendrograms. The nodes of the dendrogram are labelled with the corresponding bootstrap support values, expressed as percentages. The panels show **(A)** Illumina Hi-C data for RMV7 **(B)** Pore-C data for RMV7 **(C)** Illumina Hi-C data for RMV8 **(D)** Pore-C data for RMV8.

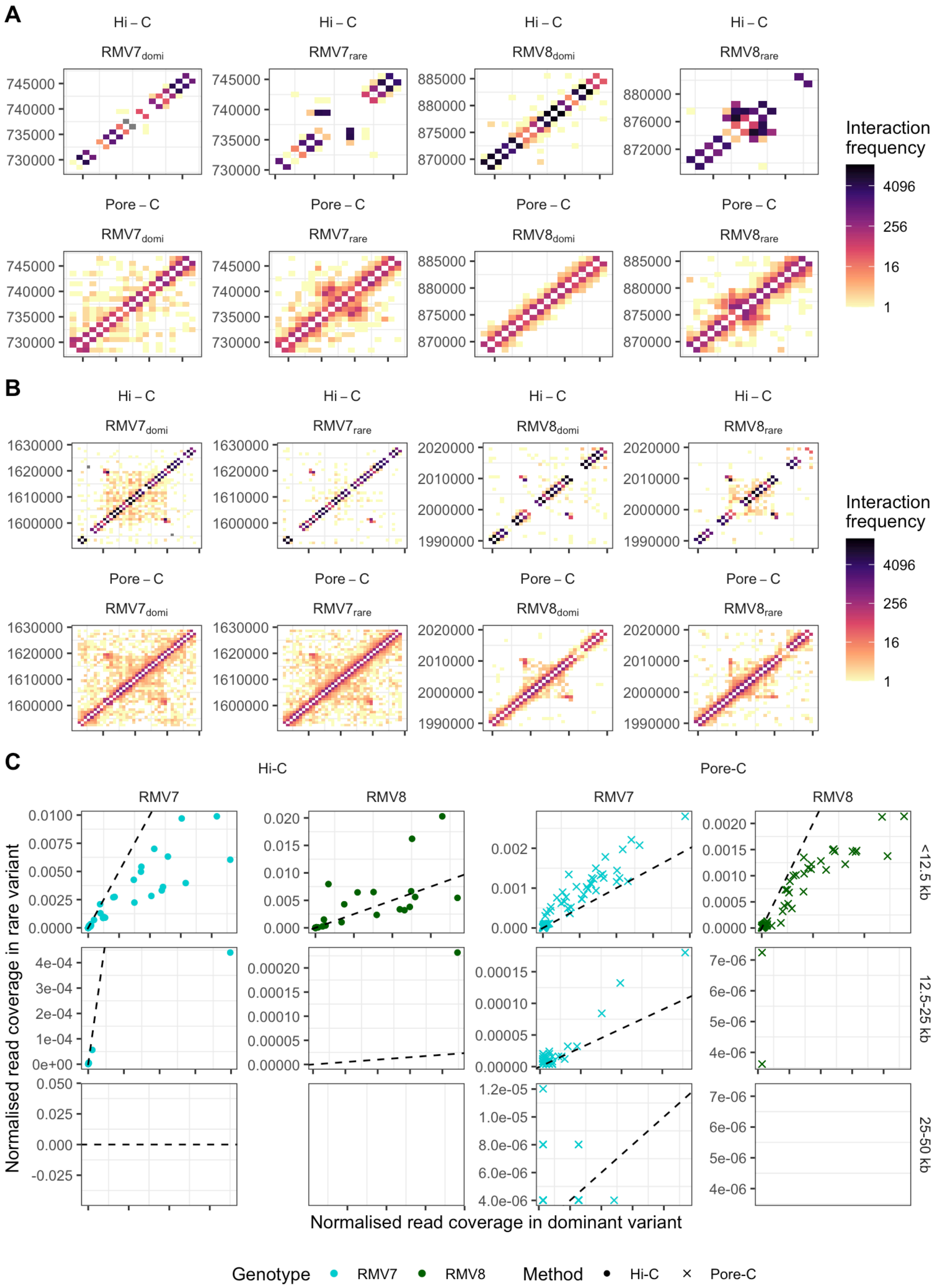


**Figure S18** Validating the differences in contact frequency in PRCIs. **(A)** Contact frequency matrices at the *tvr* loci of RMV7 and RMV8 at a resolution of 1 kb. Data are shown as in Fig. 1D. Each rare variant dataset shows distinctive off-diagonal contacts that are diagnostic of rearrangements within the *tvr* loci, relative to the dominant variant reference genome. This confirms that the dominant and rare variant datasets have been correctly identified. **(B)** Contact frequency matrices within PRCI$_{dnaN}$, for the RMV7 datasets, or within PRCI$_{malA}$, for the RMV8 datasets, at a resolution of 1 kb. Data are shown as in Fig. 1D. **(C)** Scatterplots comparing the contact frequencies within PRCI$_{dnaN}$ and PRCI$_{malA}$ between the epigenetically-distinct variants. The cumulative contact frequencies between each 1 kb locus were calculated across the three biological replicates for each combination of methodology and variant. These values were normalised by dividing them by the total number of contacts inferred in the corresponding datasets. These normalised contact frequencies were compared between the dominant and rare variants for the loci within PRCI$_{dnaN}$, in RMV7, and PRCI$_{malA}$, in RMV8. The comparisons were segregated by the distance between the interacting loci. The dashed lines are the lines of identity. These simple comparisons concur with the diffHiC and multiHiCcomparison analyses that found the Hi-C and Pore-C data implied opposing changes in the frequencies of contacts within these PRCIs between the pairs of variants.

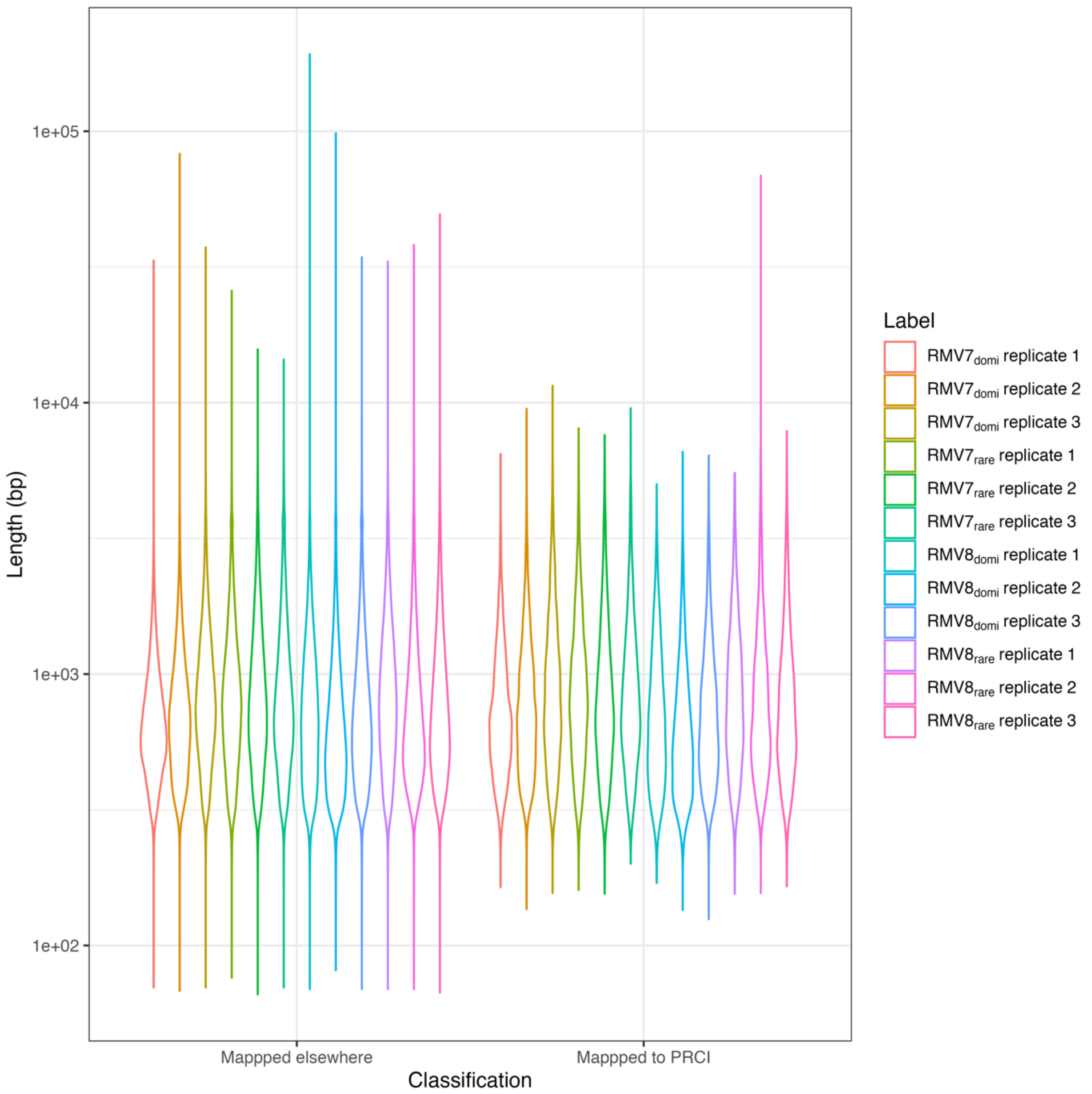


**Figure S19** Violin plots showing the distribution of Pore-C read lengths, categorised by whether or not a fragment mapped to a differentially-active PRCI: either PRCI$_{dnaN}$ (in RMV7) or PRCI$_{malA}$ (in RMV8).

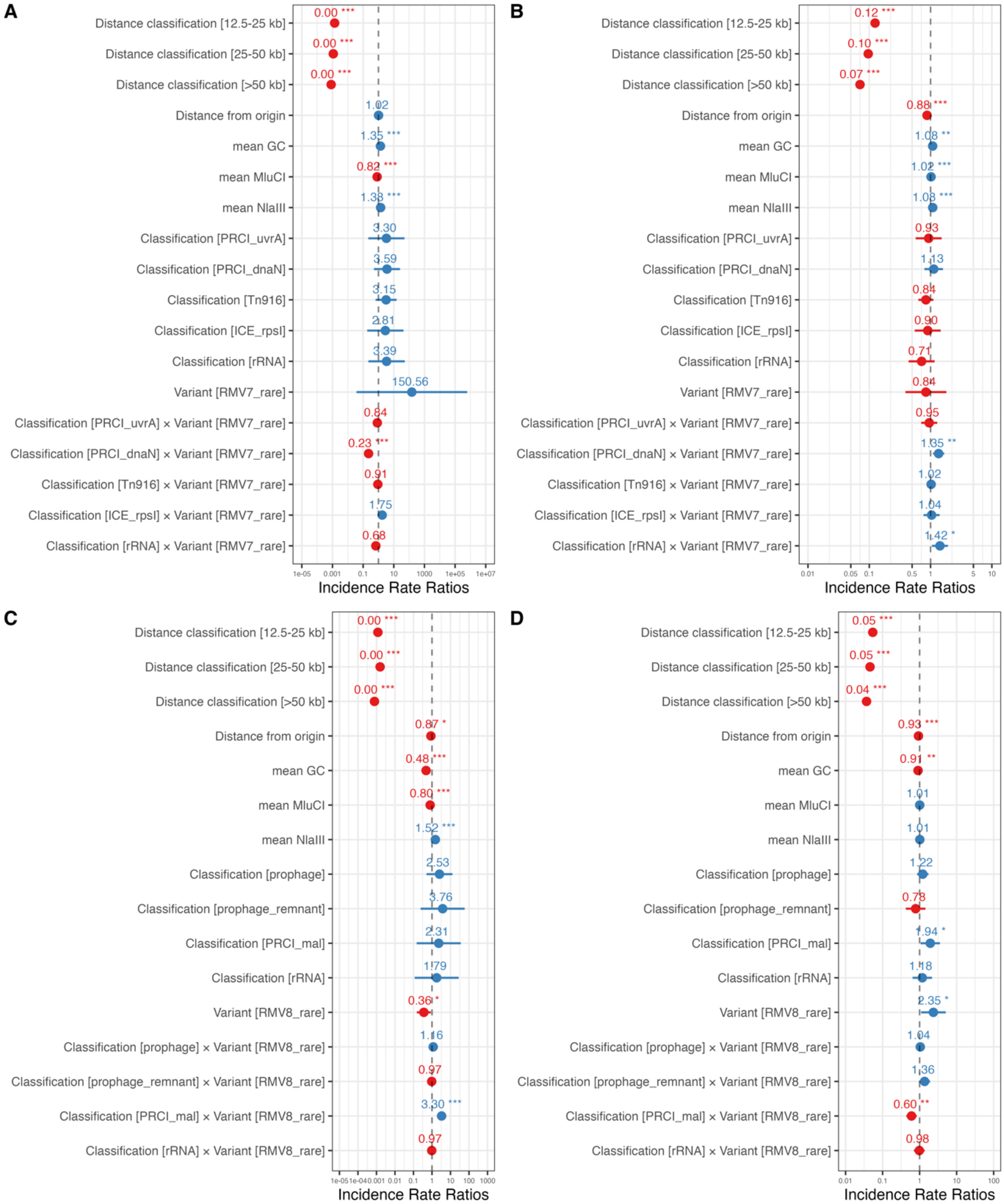


**Figure S20** Forest plots summarising the fixed effect coefficient estimates from fitting the generalised linear mixed effects models to contact matrices calculated at a resolution of 10 kb. Points represent the maximum likelihood estimate of parameter values, and are coloured by whether they increase (blue) or decrease (red) contact frequencies. The error bars represent the 95% confidence intervals of the estimates. The asterisks indicate statistically significant deviations from one: * denotes $p < 0.05$; ** denotes $p < 0.01$, and *** denotes $p < 0.001$. **(A)** Model fit to the RMV7 Hi-C data. **(B)** Model fit to the RMV7 Pore-C data. **(C)** Model fit to the RMV8 Hi-C data. **(D)** Model fit to the RMV8 Pore-C data.

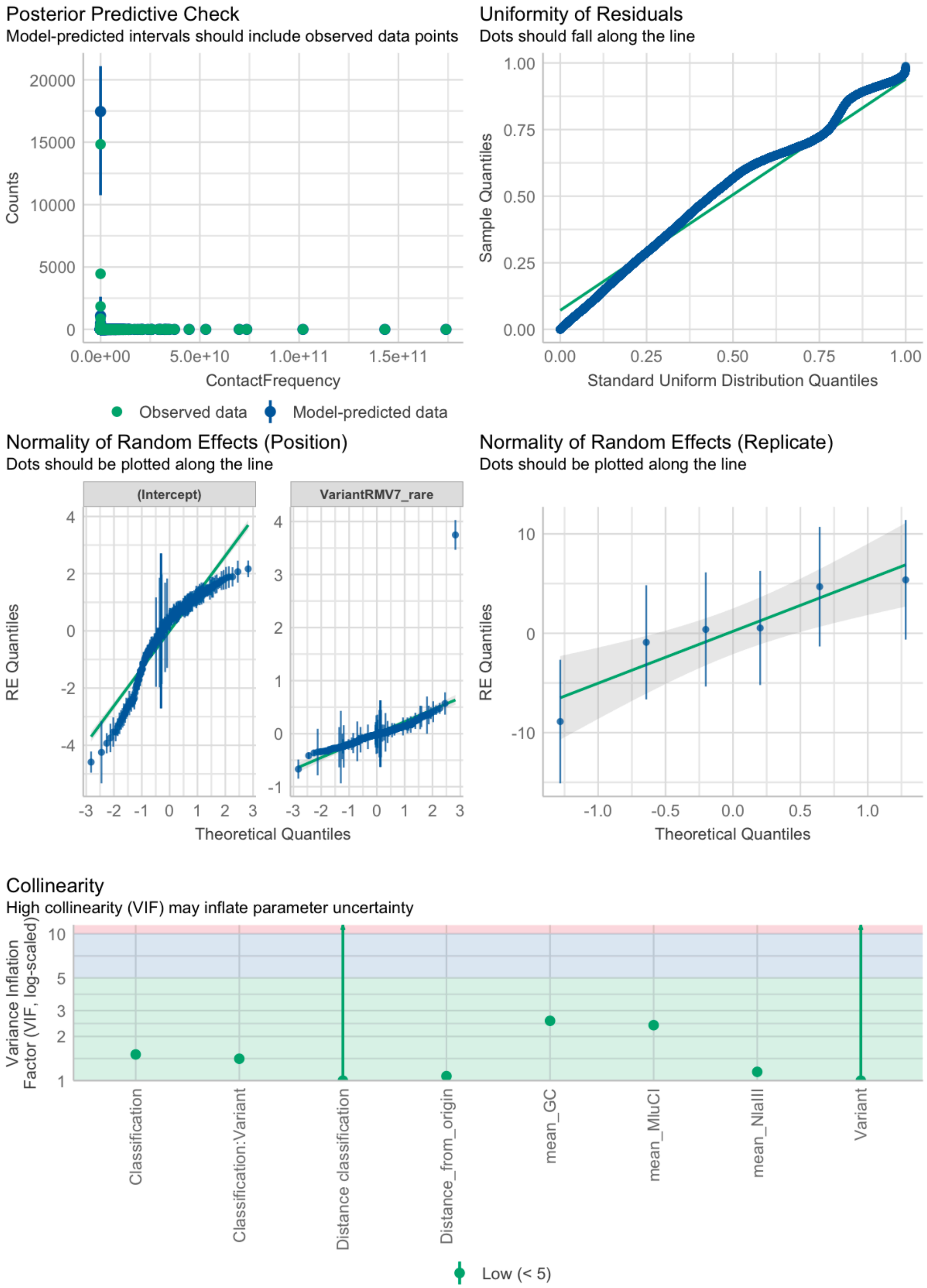


**Figure S21** Evaluation of the quality of the generalised linear mixed effects model fits to the RMV7 Hi-C data. These plots, generated by the R package performance, test whether the model performs poorly in reproducing observed values, whether it is mis-specified, or whether there are difficulties in estimating individual parameter values.

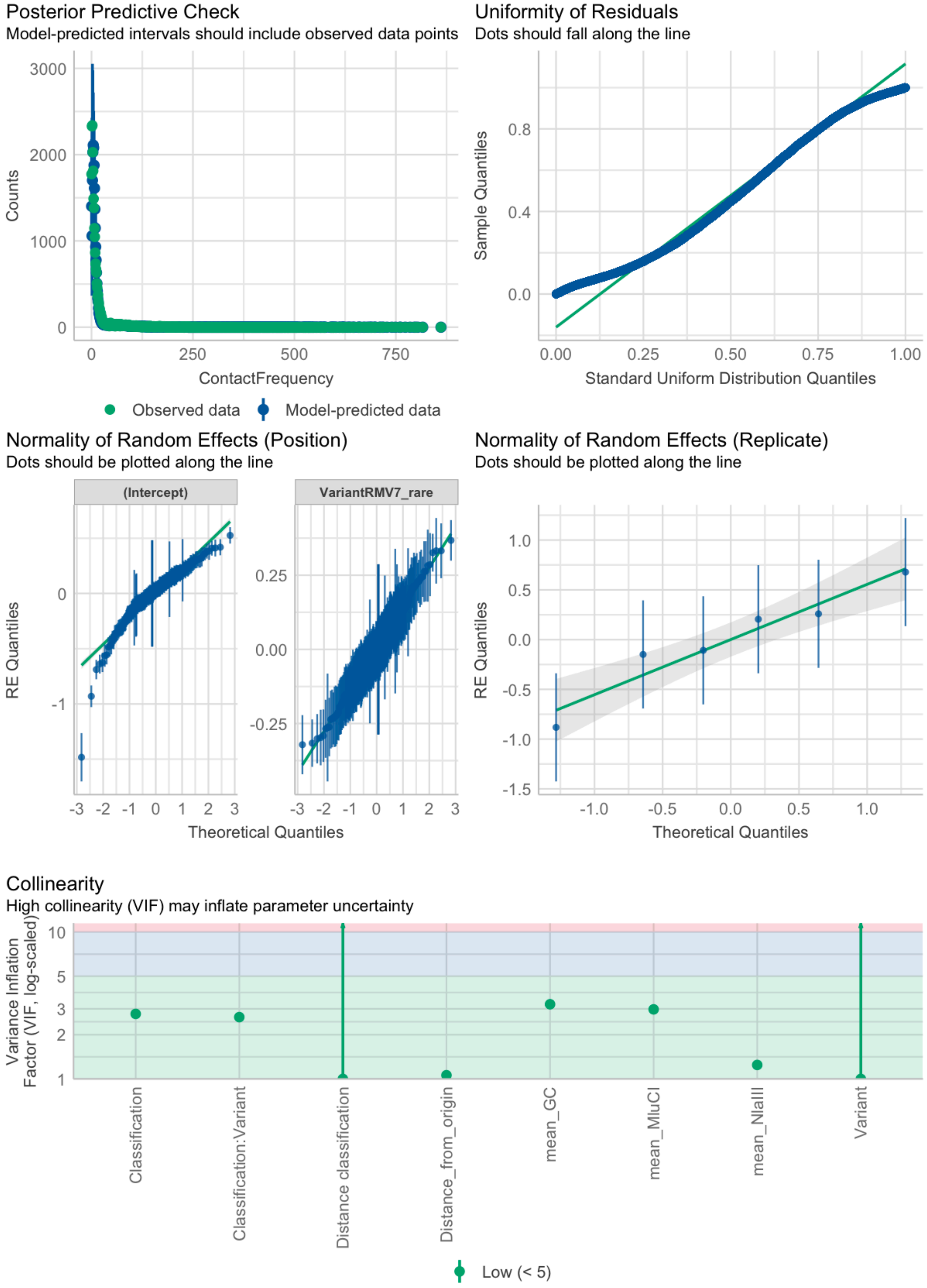


**Figure S22** Evaluation of the quality of the generalised linear mixed effects model fits to the RMV7 Pore-C data. Data are shown as in Fig. S21.

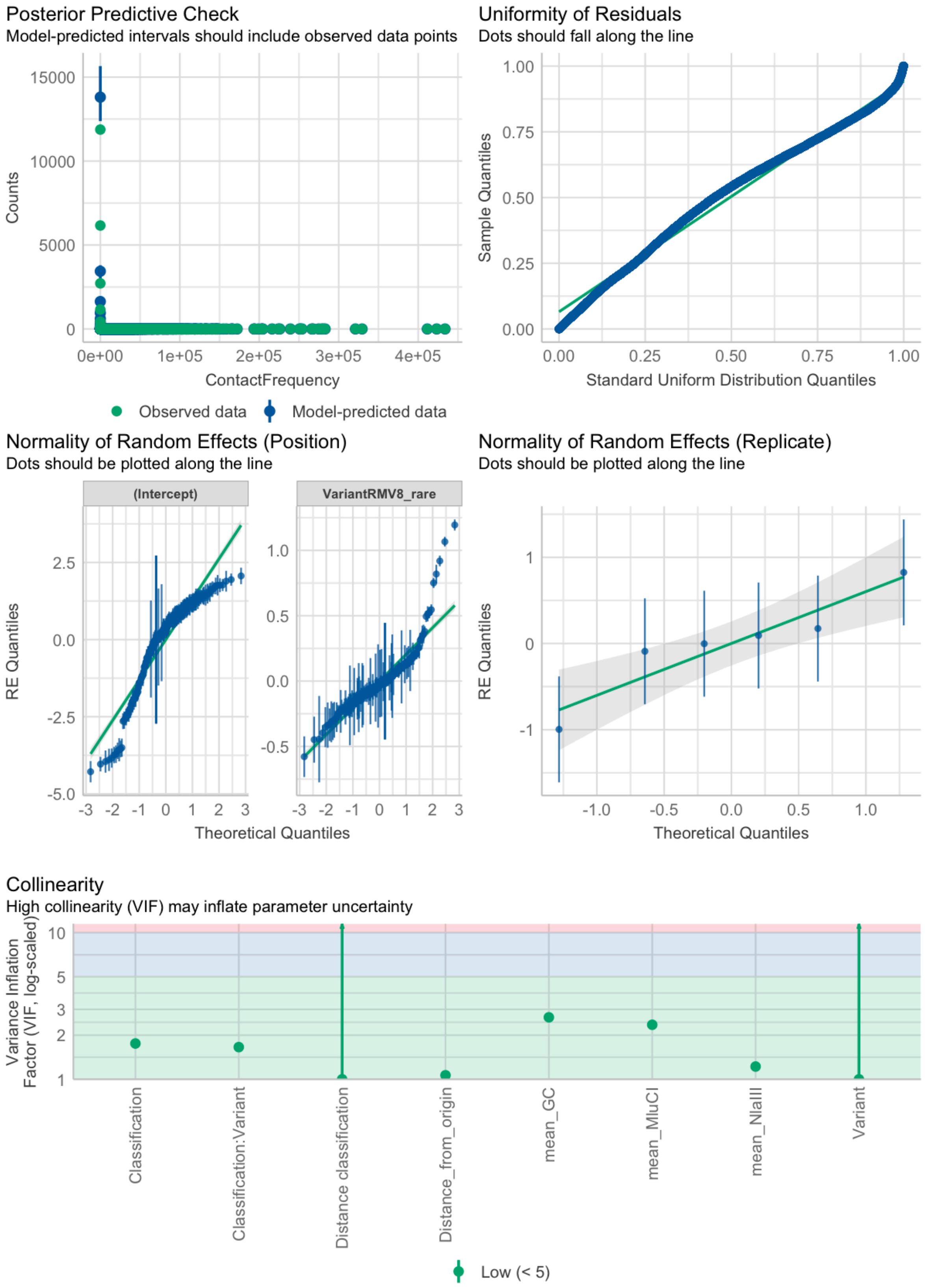


**Figure S23** Evaluation of the quality of the generalised linear mixed effects model fits to the RMV8 Hi-C data. Data are shown as in Fig. S21.

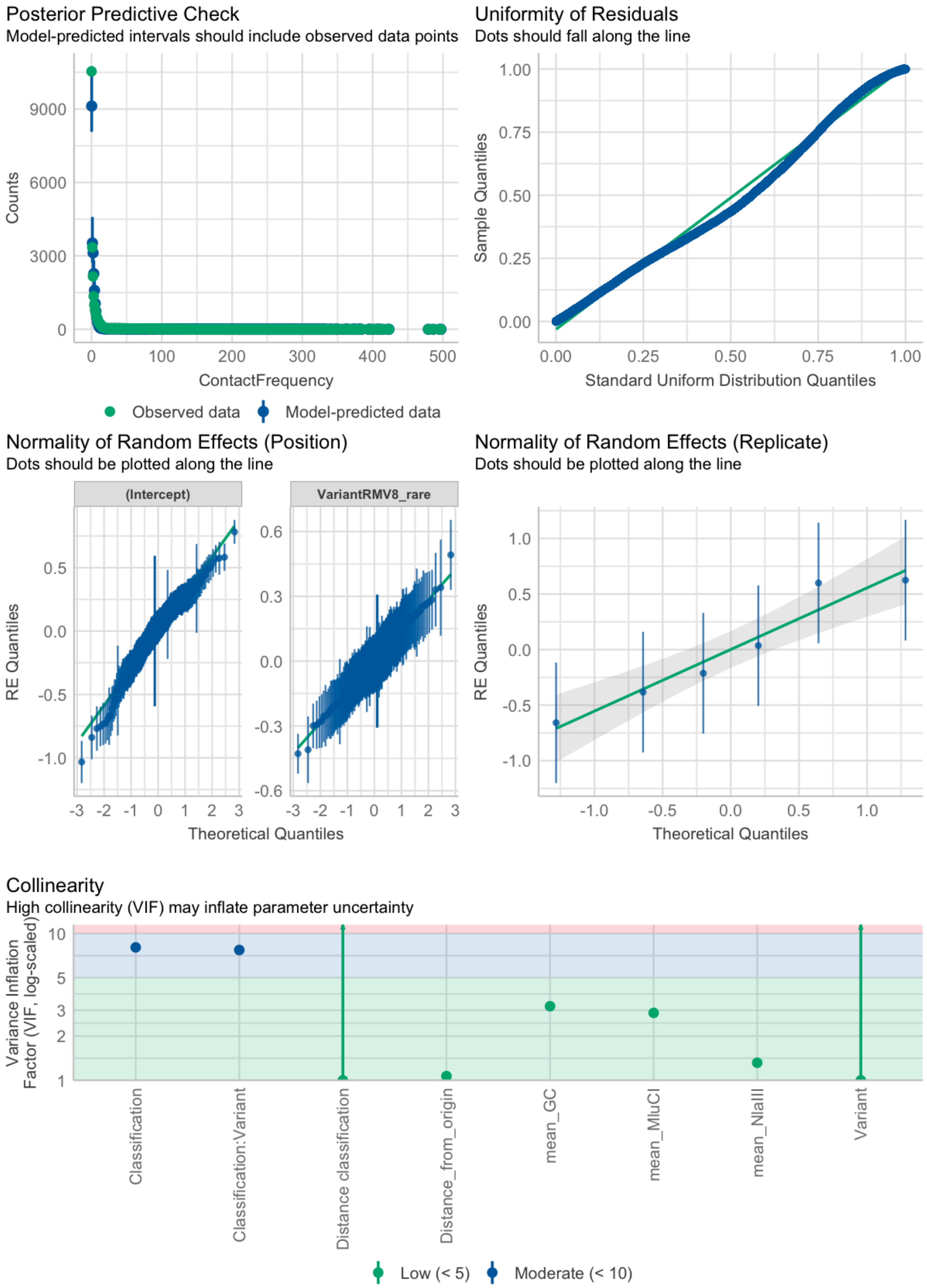


**Figure S24** Evaluation of the quality of the generalised linear mixed effects model fits to the RMV8 Pore-C data. Data are shown as in Fig. S21.

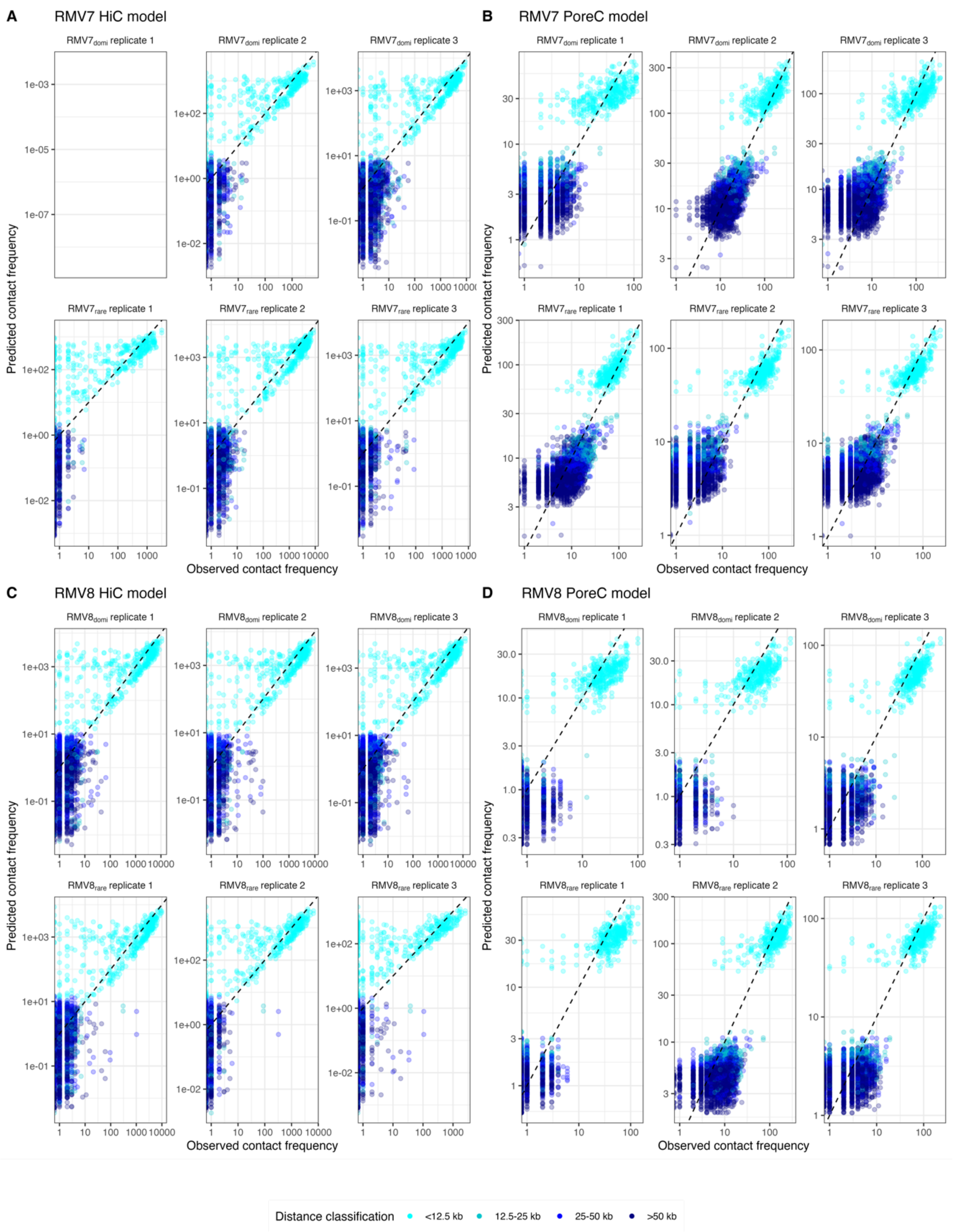


**Figure S25** Comparison of observed data with model predictions. For each model, the six panels show the comparisons between the observed and predicted contact frequencies for each locus across the biological replicates. Points are coloured by the distance between the interacting loci. Models were not fitted to the first replicate *S. pneumoniae* RMV7$_{domi}$ Hi-C dataset, which contained relatively little data. **(A)** Model predictions using the RMV7 Hi-C data. **(B)** Model predictions using the RMV7 Pore-C data. **(C)** Model predictions using the RMV8 Hi-C data. **(D)** Model predictions using the RMV8 Pore-C data.

### Supplementary Tables

**Table S1** Genotypes used in this study.

| Genotype | Description | Accession code of reference genome |
| --- | --- | --- |
| RMV7$_{domi}$ | The more common "domi" variant of RMV7 expressed a *Spn*IV allele that methylated the motif TGAN$_7$TCC. | OV904788 |
| RMV7$_{rare}$ | The less common "rare" variant of RMV7 expressed a *Spn*IV allele that methylated the motif TGAN$_7$TATC. | - |
| RMV7$_{rare}$ PRCI$_{dnaN}$::Janus | RMV7$_{rare}$ with PRCI integrated next to the gene *dnaN* replaced by a Janus cassette. | - |
| RMV8$_{domi}$ | The more common "dominant" variant of RMV8 expressed a *Spn*IV allele that methylated the motif GATAN$_6$RTC. | OX244288 |
| RMV8$_{rare}$ | The less common "rare" variant of RMV8 expressed a *Spn*IV allele that methylated the motif GTAYN$_6$TGA. | - |
| RMV8$_{rare}$ PRCI$_{malA}$::Janus | RMV8$_{rare}$ with PRCI integrated next to *malA* replaced by a Janus cassette. | - |
| RMV8$_{rare}$ ϕRMV8*::cat* | RMV8$_{rare}$ with ϕRMV8 replaced by a chloramphenicol resistance marker. | - |
| RMV8$_{rare}$ ϕRMV8*::cat* PRCI$_{malA}$::Janus | RMV8$_{rare}$ with ϕRMV8 replaced by a chloramphenicol resistance marker and PRCI integrated next to the gene *malA* replaced by a Janus cassette. | - |

**Table S2** Accession codes for Hi-C and Pore-C sequence data.

| Genotype | Replicate | Illumina Hi-C Accession Code | Nanopore Pore-C Run 1 Accession Code | Nanopore Pore-C Run 2 Accession Code |
|---|---|---|---|---|
| $RMV7_{domi}$ | 1 | ERS17696384 | ERR13946717 | ERR13946729 |
| $RMV7_{domi}$ | 2 | ERS17696392 | ERR13946718 | ERR13946730 |
| $RMV7_{domi}$ | 3 | ERS17696382 | ERR13946719 | ERR13946731 |
| $RMV7_{rare}$ | 1 | ERS17696383 | ERR13946720 | ERR13946732 |
| $RMV7_{rare}$ | 2 | ERS17696385 | ERR13946721 | ERR13946733 |
| $RMV7_{rare}$ | 3 | ERS17696386 | ERR13946722 | ERR13946734 |
| $RMV8_{domi}$ | 1 | ERS17696387 | ERR13946723 | ERR13946735 |
| $RMV8_{domi}$ | 2 | ERS17696388 | ERR13946724 | ERR13946736 |
| $RMV8_{domi}$ | 3 | ERS17696389 | ERR13946725 | ERR13946737 |
| $RMV8_{rare}$ | 1 | ERS17696390 | ERR13946726 | ERR13946738 |
| $RMV8_{rare}$ | 2 | ERS17696391 | ERR13946727 | ERR13946739 |
| $RMV8_{rare}$ | 3 | ERS17696393 | ERR13946728 | ERR13946740 |

**Table S3** Number of restriction sites in the genomes of *S. pneumoniae* RMV7$_{\text{domi}}$ and RMV8$_{\text{domi}}$, and PRCI$_{\textit{dnaN}}$ (within the RMV7$_{\text{domi}}$ genome) and PRCI$_{\textit{malA}}$ (within the RMV8$_{\text{domi}}$ genome).

| **Genome** | ***Nla*III sites** | ***Mlu*CI sites** |
|---|---|---|
| RMV7 | 6776 | 15897 |
| RMV8 | 6831 | 16425 |
| PRCI$_{\textit{dnaN}}$ | 39 | 94 |
| PRCI$_{\textit{malA}}$ | 25 | 90 |

**Table S4** Oligonucleotides used in this study.

| Name | Sequence | Function |
|---|---|---|
| RMV8_prophage_purA_chr_F (A)<br>RMV8_prophage_purA_int_F (C) | GGTCCTGGTCGTGAACAAAC<br>GCACCCTCCAAAAGCATTGA | Primers for assessing the topology of ϕRMV8. |
| RMV8_prophage_purA_chr_R (D)<br>RMV8_prophage_purA_int_R (B) | CTCAGCCTCTCTCAAAGCCT<br>AGTCCCCAAAAGCCTGAAAT | Primers for assessing the topology of ϕRMV8. |
| RMV8_A_chr_sugar_PRCI_F<br>RMV8_C_int_sugar_PRCI_F | ATTTATCCCCGCCACCCTTT<br>AGCCACTTATCCAAAGACAACA | Primers for assessing the topology of $PRCI_{malA}$. |
| RMV8_B_int_sugar_PRCI_R<br>RMV8_D_chr_sugar_PRCI_R | TGAGGTGTGAAATGATGCCT<br>GGAAAATGGATATGGAAGCATAGGT | Primers for assessing the topology of $PRCI_{malA}$. |
| RMV7_PRCI_uvrA_Chr_A_F<br>RMV7_PRCI_uvrA_int_C_F | GGTTCAAGCGTATCGATCTCT<br>GCGCGTGATATAAGGGTTAGG | Primers for assessing the topology of $PRCI_{uvrA}$. |
| RMV7_PRCI_uvrA_int_B_R<br>RMV7_PRCI_uvrA_chr_D_R | TCTACGAAACTGCACTAAAAGC<br>GCGCCCCATGAATGACAATT | Primers for assessing the topology of $PRCI_{uvrA}$. |
| RMV7_A_For_Chr<br>RMV7_C_For_int | AAAAGGCTAATCGTTGGGAAATT<br>GGCTCGGTCATGCAAAACTT | Primers for assessing the topology of $PRCI_{dnaN}$. |
| RMV7_B_Rev_int<br>RMV7_D_Rev_Chr | AATATCAAGGGTTTAGGCGCT<br>CCAACGATACCTGCTGTCA | Primers for assessing the topology of $PRCI_{dnaN}$. |
| malA_KO_UP_For<br>malA_KO_UP_Rev_ApaI<br>malA_KO_DOWN_For_BamHI<br>malA_KO_DOWN_Rev | CAAAGCCTTACTTACCTTTACCTGAT<br>AAGGGCCCGAGTATAAAATGAAAAAGG<br>AAGGATCCCCTAGACTTGAAATAAAGC<br>TGAATAACACGGGTTACGGTTGAAG | Primers used to generate amplicons for the replacement of $PRCI_{malA}$ with a Janus cassette. |
| rpoA_Rev<br>rpoA_For | CACGAGCAGGTTCCACTTGA<br>TGGTCGTGGATATGTACCTGC | Primers for the amplification of *rpoA* as a standard for qPCR experiments. |
| Janus_For<br>Janus_Rev | TTGGGCCCCCGTTTGATTTTTAATGGATAATGTG<br>ATGGATCCCCTTTCCTTATGCTTTTGGACG | Primers for the amplification of the Janus cassette for mutant construction. |
| sugar_RMV8_qRT_For_int_tn2<br>sugar_RMV8_qRT_Rev_int_tn2 | GCCCGTGATATTATCGTGCC<br>GAATACGCAAACCAGTCGCT | Primers for the quantification of $PRCI_{malA}$ *int* gene expression |
| qRT_dnaC_RMV8_PRCI_For<br>qRT_dnaC_RMV8_PRCI_Rev | TACAACCATTTTGCCCGGAG<br>GGTATTTCTGAGCTTGCCCC | Primers for the quantification of $PRCI_{malA}$ *dnaC* gene expression |
